# Connecting secretome to hematopoietic stem cell phenotype shifts in an engineered bone marrow niche

**DOI:** 10.1101/2020.01.19.911800

**Authors:** Aidan E. Gilchrist, Brendan A.C. Harley

## Abstract

Hematopoietic stem cells (HSCs) primarily reside in the bone marrow, where they receive external cues from their local microenvironment. The complex milieu of biophysical cues, cellular components, and cell-secreted factors regulates the process by which HSC produce the blood and immune system. We previously showed direct co-culture of primary murine hematopoietic stem and progenitor cells with a population of marrow-derived mesenchymal stromal and progenitor cells (MSPCs) in a methacrylamide-functionalized gelatin (GelMA) hydrogel improves hematopoietic progenitor maintenance. However, the mechanism by which MSPCs influenced HSC fate decisions remained unknown. Herein, we report the use of proteomic analysis to correlate HSC phenotype to a broad candidate pool of 200 soluble factors produced by combined mesenchymal and hematopoietic progeny. Partial Least Squares Regression (PLSR), along with an iterative filter method, identified TGFβ-1, MMP-3, c-RP, and TROY as positively correlated with HSC maintenance. Experimentally, we then observe exogenous stimulation of HSC monocultures in GelMA hydrogels with these combined cytokines increases the ratio of hematopoietic progenitors to committed progeny after a 7-day culture 7.52 ± 3.65 fold compared to non-stimulated monocultures. Findings suggest a cocktail of the downselected cytokines amplify hematopoietic maintenance potential of HSCs beyond that of MSPC-secreted factors alone. This work integrates empirical and computation methods to identify cytokine combinations to improve HSC maintenance within an engineered HSC niche, suggesting a route towards identifying feeder-free culture platforms for HSC expansion.

**Insight:** Hematopoietic stem cells within an artificial niche receive maintenance cues in the form of soluble factors from hematopoietic and mesenchymal progeny. Applying a proteomic regression analysis, we identify a reduced set of soluble factors correlated to maintenance of a hematopoietic phenotype during culture in a biomaterial model of the bone marrow niche. We identify a minimum factor cocktail that promotes hematopoietic maintenance potential in a gelatin-based culture, regardless of the presence of mesenchymal feeder-cells. By combining empirical and computational methods, we report an experimentally feasible number of factors from a large dataset, enabling exogenous integration of soluble factors into an engineered hematopoietic stem cell for enhance maintenance potential of a quiescent stem cell population.

## 1. Introduction

Hematopoietic stem cells (HSCs) maintain the body’s system of blood and immune cells via hematopoiesis, whereby a small population of cells (<0.007% of murine bone marrow) produce half a trillion cells daily (1). This process is highly regulated by the local microenvironment, termed *niche*, which provides a diverse mixture of biophysical, cellular, and soluble factor cues (2-11). The complex milieu of the bone marrow, the primary location of adult HSCs, presents orders of magnitude of moduli from 0.25 – 25 kPa (12), cellular components from stromal, hematopoietic, and nervous systems (13-15), as well as gradients in biomolecular and metabolic factors (16, 17). Together, these factors establish distinct hierarchically organized niches that maintain a homeostasis of HSC proliferation and lineage specification versus self-renewal and quiescence.

Destruction or removal of HSCs from the context of their native bone marrow niche can lead to hindered hematopoiesis and erratic differentiation (18, 19). This is of immediate concern for HSC transplants which are commonly used to treat disorders of the blood and immune system and chemo-radiation therapy (20). As such, the microenvironment of the HSC population is of both clinical and engineering relevance, demonstrating the need to engineer a culture platform that can maintain, condition, or expand HSCs prior to transplantation (21-23). Inspired by distinct niche compartments within the bone marrow that govern distinct hematopoietic activity of quiescence or activation, culture platforms that leverage the diverse range and tunability of biomaterials have been developed to engineer stem cell fate in an artificial niche (24-27). Both elasticity and stress relaxation time-scales of the biomaterial substrate have been shown to bias stem cell fate (28, 29), and decoration of covalently-bound factors simulates presentation of ECM proteins which lead to either maintenance or proliferation of a stem cell population (30-33). Additionally, the coupling of biomaterials and niche-associated cells synergistically moderate cell-cell interactions through direct cell-cell contact or soluble factors (34-36). Recent work in non-adherent liquid culture has demonstrated expansion of HSCs when presented with a cocktail of soluble factors, however this has not been translated to a biomaterial culture system where cell signaling is modulated by the matrix environment (21). Within a biomaterial, biotransport of cell-secreted factors can be externally controlled via perfusion and flow (36, 37), while electrostatic hindrance and ECM-binding motifs can be modulated for intrinsic control of soluble factor biotransport (38-42).

We and others have demonstrated the coupling of biophysical cues and cellular signals from bone-marrow derived cells induces differential hematopoietic lineage and cell-cycle patterns (43-45). We have previously reported (46) that a methacrylamide-functionalized gelatin (GelMA) hydrogel, in combination with co-cultured mesenchymal stromal and progenitor cells (MSPCs), can be used to elucidate external cues that drive a hemopoietic response. Within this system, soluble signaling from MSPCs induced a quiescent HSC state in a stiffer environment (modulus ∼10^1^ kPa) compared to a soft environment (modulus ∼10^0^ kPa). The exact factors and mechanism that drove HSC response were unidentified, inspiring efforts to understand the driving forces within a GelMA culture platform. As soluble signaling dominated cell-cell interactions, additional secretome analysis is needed to elucidate the biomolecular factors involved in modulating hematopoietic activity.

The objective of this study is to identify soluble factors that lead to higher HSC maintenance in an artificial stem cell culture. We used a statistical framework, Partial Least Squares Regression (PLSR), to correlate secretome information to hematopoietic differentiation patterns and downselect to factors with the greatest impact on hematopoietic fate. We subsequently examined the response of hematopoietic stem and progenitor cells (HSPCs) to exogenous stimulation by a subset of cytokines identified by the model. HSPCs were stimulated in single-culture or in the presence of MSPCs to define indirect or direct effects of cytokines that were not accounted for in the PLSR model. Taken together, we have demonstrated the use of secretome analysis to identify a subset of soluble factors that can be used to increase the maintenance potential of an HSPC culture platform without the need for feeder-cell signaling.

## 2. Materials and Methods

### 2.1 Quantitative measurement of soluble factors

#### 2.1.1 Hematopoietic and mesenchymal cell isolation

All work involving primary cell extraction was conducted under approved animal welfare protocols (Institutional Animal Care and Use Committee, University of Illinois at Urbana-Champaign). Murine hematopoietic stem and progenitor cells (HSPCs) and mesenchymal stromal progenitor cells (MSPCs) were cultured in methacrylamide-functionalized gelatin (GelMA) following the isolation and culture protocols outlined previously (46). In brief, HSPCs were isolated from the crushed tibia and femur of C57BL/6 female mice, age 4 – 8 weeks (The Jackson Laboratory). Initial hematopoietic lineage negative enrichment was performed with EasySep™ Mouse Hematopoietic Progenitor Cell Isolation Kit (#19856, Stemcell Technologies, CA), followed by collection of the Lin^-^ Sca-1^+^ c-kit^+^ (LSK) fraction using a BD FACS Aria II flow cytometer. LSK antibodies were supplied by eBioscience (San Diego, CA), and are as follows: APC-efluor780-conjugated c-kit (1:160, #47-1172-81), PE-conjugated Sca-1 (0.3:100, #12-5981-83), and Lin: FITC-conjugated CD5, B220, CD8a, CD11b (1:100, #11-0051-82, #11-0452-82, #11-0081-82, #11-0112-82), Gr-1 (1:400, #11-5931-82), and Ter-119 (1:200, #11-5921-82) (1, 47, 48). MSPCs were isolated from the crushed bone following a commercially available protocol and cultured for 10 days prior to collection and use (#05513, Stemcell Technologies).

#### 2.1.2 Hematopoietic and mesenchymal conditioned media

HSPCs and MSPCs were cultured at increasing ratios of 1:0, 1:1, 1:10, 0:1, 0:10 (HSPCs:MSPCs) in 4, 5, 7.5 wt% methacrylamide-functionalized gelatin (GelMA) (49, 50) at constant crosslinking conditions (85% functionalization, 0.1% lithium acylphosphinate photoinitiator (PI), and 7.14 mW/cm^2^ UV light for 30 seconds) for a total of 15 conditions (Figure 1A) (33, 46, 51). HSPC seeding density was kept constant at 1×10^5^ HSPCs/mL, and cells were encapsulated in 5 mm diameter hydrogels (20 µL) and cultured for 7 days in 300 µL SFEM media (#09650 Stemcell Technologies) supplemented with 100 ng/mL SCF (#250-03, Peprotech) and 0.1% PenStrep, changed every 2-days. Media was collected from each sample at day 2, 4, and 6 and stored at −80°C for use in secretome analysis. Hematopoietic differentiation patterns in response to hydrogel and seeding condition were previously reported by Gilchrist et al. and are publicly available (46). Importantly, the collected media and hematopoietic lineage patterns were from the same samples, allowing for direct mapping of sample conditioned media to HSC phenotype.

**Figure 1.**
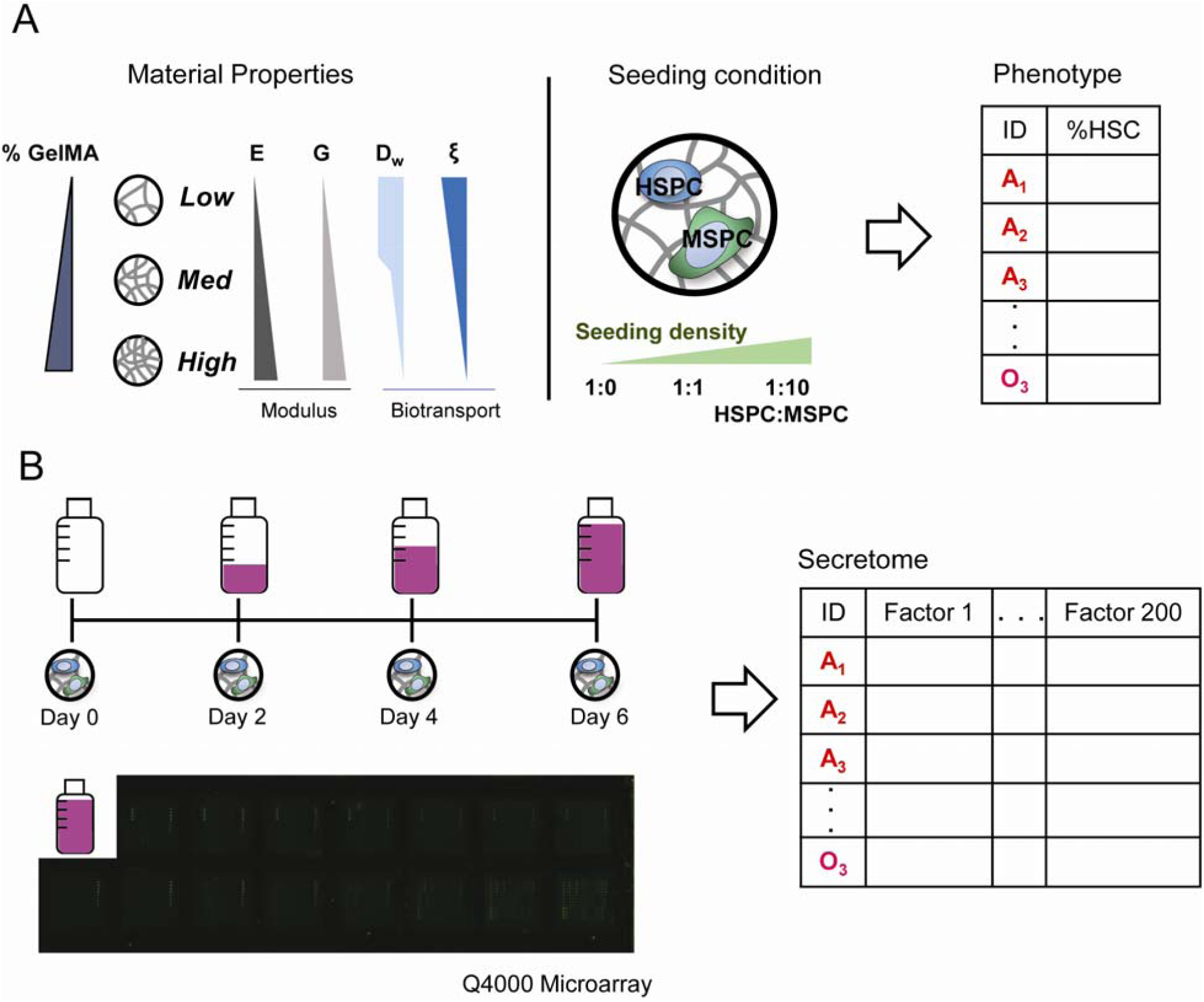
Overview of cytokine data collection. **A**. A library of GelMA hydrogels were produced with disparate properties (E: elastic modulus, G: shear modulus, D_w_: diffusion of water, ξ: mesh size). Within the *Low, Med*, and *High* hydrogels, hematopoietic stem and progenitor cells (HSPC) and mesenchymal stem progenitor cells (MSPC) were co-cultured at seeding ratios of 1:0, 1:1, and 1:10 HSPC:MSPC. **B.** Media was changed and collected at days 2, 4, and 6 and pooled together before analysis via a quantitative cytokine microarray analysis.

#### 2.1.3 Soluble factor quantification in conditioned media

Media collected from days 2, 4, and 6 from each sample was pooled (sample A_1_: day 2, 4, 6) and used for subsequent proteomic analysis: 15 conditions, in triplicate (Figure 1B). A panel of 200 murine cytokines was quantitatively assessed via a Quantibody cytokine array (#QAM-CAA-4000-1, RayBiotech, Norcross, GA). The full list of cytokines can be found on the manufacturer’s website along with a detailed protocol and, for ease of access, has been provided in Supp. Table 1. In brief, each well of the microarray was blocked and washed prior to incubation with 100 µL of sample overnight. The wells were then washed, incubated with a biotinylated antibody cocktail, and then incubated with a Cy3 equivalent dye-streptavidin. Each microarray also included a standard curve dilution for later quantitative analysis. The microarrays were sent to RayBiotech for imaging. Quantitative results were extracted from the raw data using Q-analyzer software (Analysis Tool for QAM-CAA-4000, RayBiotech) with concentration values set within limits of detection.

**Table 1.** Glass effect size (Glass *Δ) is shown with 1:0 None as the control group. This is a measure of the effect a cytokine has on Progenitor Factor compared to the unstimulated HSPC-only condition. Sample size for each condition is listed under effect size. L: latent MMP3. A: activated MMP3.

| HSPC:MSPC ratio | TGFβ-1 | MMP3 (L) | MMP3 (A) | TROY | cRP | Combo (L) | Combo (A) | None |
| --- | --- | --- | --- | --- | --- | --- | --- | --- |
| 1:0 | 25.9<br>n=10 | 16.6<br>n=7 | -5.88<br>n=6 | 27.5<br>n=8 | 9.55<br>n=8 | 31.6<br>n=10 | 3.51<br>n=6 | 0<br>n=16 |
| 1:1 | 21.6<br>n=6 | -3.60<br>n=4 | -3.17<br>n=6 | 2.31<br>n=4 | -5.43<br>n=4 | 35.1<br>n=6 | -2.90<br>n=6 | -1.28<br>n=12 |

### 2.2 Iterative filter method

#### 2.2.1 Pre-processing

Partial Least Squares Regression (PLSR) requires a y-variable by which we can correlate x-variables, as such, only conditions with hematopoietic lineage data were used for regression analysis (*Low, Med, High*; 1:0, 1:1, 1:10 HSPCs:MSPCs). From these conditions, the y-variable was taken as the proportion of HSCs (CD34^+/-^ CD135^-^ LSK) in the hematopoietic population (LSK, Common Myeloid Progenitors, and lineage committed), after a 7-day culture, while the predictors (x-variables) for the HSC response are the concentrations of the 200 cytokines. As there are potential interaction effects from cytokines that impact HSCs response, the predictors were Log transformed. As some samples had cytokine concentrations of zero or below the limit of detection, care was taken in Log transform to deal with undefined values. Broadly shifting the data by a constant factor, ε, can lead to variable results depending upon the value chosen (52, 53), therefore, we chose to add a variable shift factor, reflective of the popular commercially available SIMCA package (54):

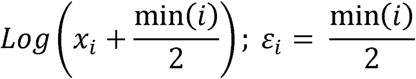

Where *i* refers to cytokine *i*, and *ε*_*i*_ is equal to half the minimum cytokine concentration; if all samples had non-zero values for cytokine *i* then *ε*_*i*_ was equal to zero. The transformed data was then centered around zero and scaled to unit variance (55, 56).

#### 2.2.2 Identification of cytokines for HSC response

Cytokine concentration was correlated to the HSC proportion, via PLSR using the mixOmics package in RStudio (57, 58). The independent x-block consisted of 200 columns, with 27 rows corresponding to 9 culture conditions in triplicate. The dependent block was a single variable of HSC proportion with 27 rows. The inner iterative process ran PLSR on an initially full dataset (f), and 10-fold cross-validation was performed, with 50 repetitions. The number of components in the model was determined by a cross-validated metric of predictive power, Q^2^, of less than 0.05. The variable importance projection (VIP) was then calculated for each cytokine, and the lowest scoring cytokine was removed. PLSR was then repeated on this f-1 dataset. This iteration was repeated, with storing of model metrics and removal of min(VIP) cytokine, until a model could no longer be constructed. The optimal model from the iteration process was then defined as the model with a Q^2^ within 1 standard error of the mean from the maximum occurring Q^2^ across all the models (59). This entire process was then repeated (n=1000), and cytokines appearing in more than 98% of the optimized models were compile into a final reduced model. A cutoff of 98% was chosen to limit the number of cytokines to a strict significant level, analogous to α=0.02.

#### 2.2.3 Identify autocrine or paracrine signaling molecules

Cytokine concentrations of the reduced model were compared across single cultures of HSPCs and MSPCs (HSPC:MSPC; 1:0, 0:1). Significance was determined with Welch’s ANOVA with Tukey’s pairwise means comparison. Significance was examined across seeding condition (1:0 and 0:1) and the 7.5 wt% GelMA hydrogel (*High*; 1:0, 0:1).

#### 2.2.4 Data visualization

Heatmap and principal component analysis (PCA) was performed in Matlab (Natick, MA). For visualization of the entire microarray, the dataset was centered and scaled to unit variance. Visualization of the full and reduced model for PLSR used the transformed, scaled, and centered data. All clustering was performed with average Euclidean distance (60).

### 2.3 Model validation and stimulation of stem cell culture

#### 2.3.1 Exogenous stimulation

Freshly harvested HSPCs and MSPCs were encapsulated in 20 µL of 7.5 wt% GelMA hydrogel (DOF=85%, PI=0.1%) as described previously, at a seeding density of 1×10^5^ HSPCs/mL and ratio of 1:0 or 1:1 HSPCs:MSPCs (46). Each cell-hydrogel construct was placed in individual wells (48-well plate) and were cultured for 7 days. Cells were maintained in SFEM media supplemented with 100 ng/mL SCF and 0.1% P.S, and 50 ng/mL of the cytokines identified by the reduced model. Cytokines were added either individually, a combination of all the cytokines, or none of the cytokines: recombinant murine TGFβ-1 (#763104, BioLegend), recombinant murine MMP-3 (#552704, BioLegend), recombinant murine c-RP (#50409-M08H, Sino Biological), recombinant murine TROY (#50148-M08H, Sino Biological). MMP-3 was activated in 0.1 mM P-Aminophenylmercuric acid (#164610, Milipore Sigma) at 37 °C for 24 hours. Media changes occurred every two days.

#### 2.3.2 Cell lineage and cycle analysis

To protect against cell loss during the fixation and wash steps, two individually-cultured hydrogels were pooled for each sample. Sample dissociation was performed in 500 µL of PBS + 25% FBS and 100 Units Collagenase Type IV (#LS004186, Worthington Biochemical). Samples were made piecemeal using scissors, and then placed on a rotator (200 rpm) at 37°C for 30 minutes. Degradation was quenched with 1 mL PBS + 5% FBS and centrifuged at 300 rcf x 10 minutes. The collected pellet was resuspended in PBS + 5% FBS and stained with surface marker antibodies. Following staining, the cells were fixed and permeabilized with Foxp3/Transcription Factor Staining Buffer Set (#00-5523-00, ThermoFisher), and then stained with the intranuclear Ki-67 stain and DAPI (1mg/mL, 10:300, #D21490, ThermoFisher). Cells were resuspended in PBS + 5% FBS and analyzed via Fluorescence-Assisted Cytometry (FACs), using a BD LSR Fortessa (BD Biosciences, San Jose, CA). Lysed whole bone marrow was used to create fluorescent minus one (FMO) controls for gating. DAPI was used to discriminate cells from debris, and analysis was performed with a 5,000 DAPI parameter thresholding. Cells were classified as Long-Term repopulating HSCs (LT-HSCs: CD34^-^ CD135^-^ Lin^-^ Sca1^+^ c-kit^+^) (61-63); Short-Term repopulating HSCs (ST-HSCs: CD34^+^ CD135^-^ LSK) (61-63); HSCs (CD34^+/-^ CD135^-^ LSK); or Multipotent Progenitors (MPPs: CD34^+^ CD135^+^ LSK) (63, 64). Cell cycle was classified as G0 (Ki-67^-^, DAPI^≤2N^); G1 (Ki-67^+^, DAPI^≤2N^); SGM (DAPI^>2N^). All antibodies were supplied by eBioscience (San Diego, CA), and are as follows: PE-conjugated Ki-67 (0.3:100, #12-5698-82), eFluoro660-conjugated CD34 (5:100, #50-0341-82), PE-CY5-conjugated CD135 (5:100, #15-1351-82), APC-efluor780-conjugated c-kit (1:160, #47-1172-81), PE-CY7-conjugated Sca-1 (0.3:100, #25-5981-81), and Lin: FITC-conjugated CD5, B220, CD8a, CD11b (1:100, #11-0051-82, #11-0452-82, #11-0081-82, #11-0112-82), Gr-1 (1:400, #11-5931-82),and Ter-119 (1:200, #11-5921-82).

### 2.4 Statistical Analysis

Prior to significance testing via ANOVA, normality and equality of variance were tested with Shapiro-Wilks and Brown–Forsythe at significance level 0.05 (65, 66). Significance of ½ power transformed Progenitor Factor data with unequal variance was examined with Welch’s modified 2-way ANOVA and post hoc pairwise means comparisons with Dunnett T3 (67). Cell-cycle significance was tested with Kruskal-Wallis and post hoc pairwise means comparison with Dunn Test and FDR adjusted p-value. Glass’s Δ* effect size for t-test with small control group was used to measure influence of cytokines against a non-stimulated, single-culture control group (68). All statistical analysis was performed in R.

### 2.5 Data Sharing Statement

Original data and R code are available upon request from the corresponding author. All flow cytometry files have been uploaded to flowrepository.org (FR-FCM-Z2EP). All microarray data has been uploaded to NCBI Gene Expression Omnibus (<u>GSE143987</u>).

## 3. Results

### 3.1 Visualizing the entire secretome dataset

We first applied standard techniques for visualization and analysis of differentially expressed cytokines, i.e. Principal Component Analysis (PCA) and heatmap with dendrogram (Figure 2). However, this did not reveal substantial data-based inference on hematopoietic response. Notably, while we observe clustering of the single culture, low stiffness condition (A; *Low;* 4wt% GelMA, 1:0), with an increased concentration of cytokines compared to other conditions (Figure 2B), the magnitude of the data does not innately provide insight regarding secretome elements most suitable for HSC maintenance. Similarly, while the dimensionality reduction of the data (PCA) shows two “nodes” of activity (Figure 2C) there is no readily apparent biological significance to the groups as they do not cluster according to condition or hematopoietic response. A separate heatmap file has been provided for ease of legibility (Supp.Figure 1).

**Figure 2.**
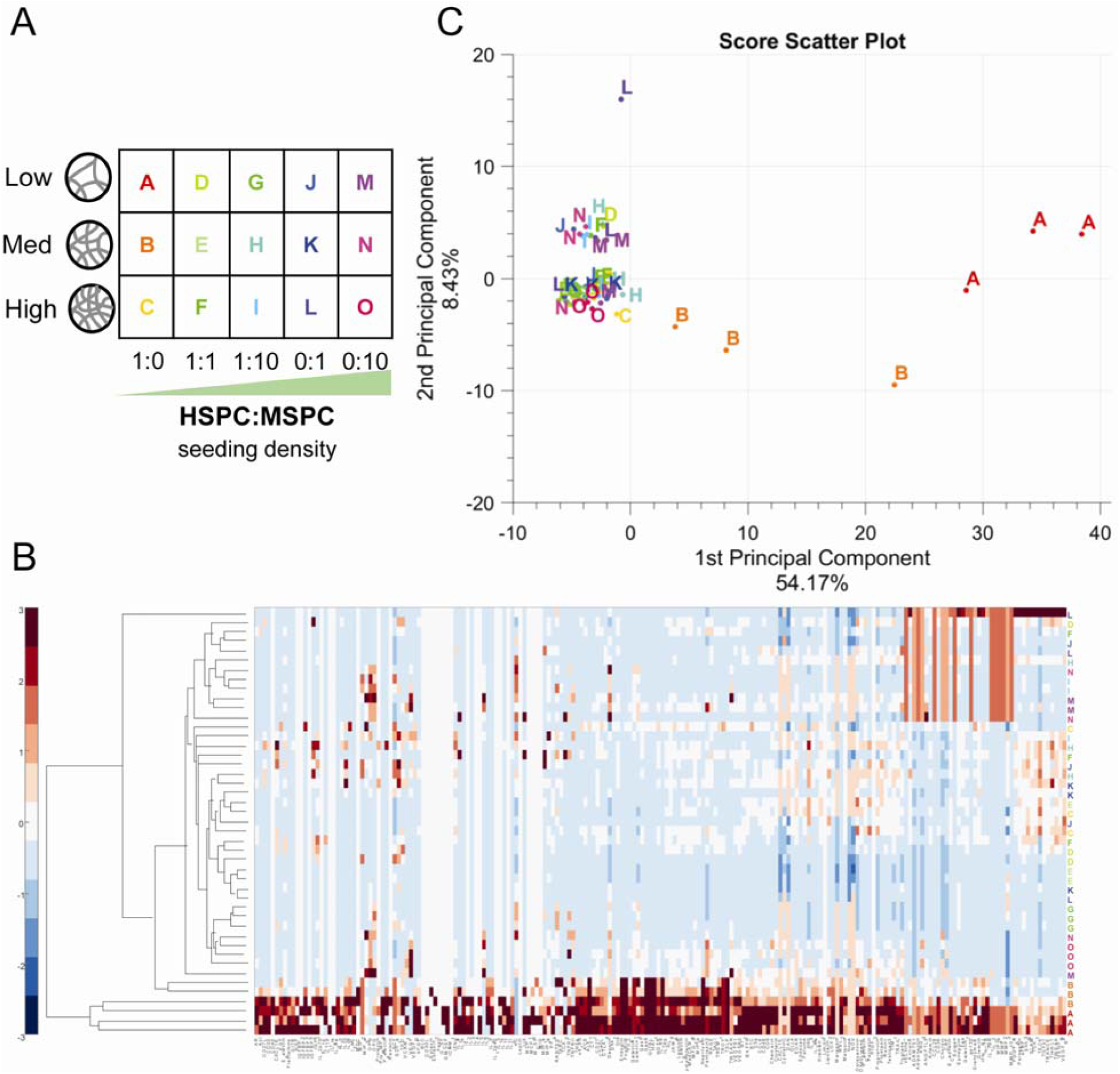
First stage analysis of complete data set. **A**. Conditions shown in visualization of dataset. **B**. Heatmap of cytokines across all conditions. The data is centered and scaled along the columns (cytokines). **C**. Principal component analysis of all conditions (centered and scaled), showing the first two components.

### 3.2 Model-based reduction of the secretome data

To downselect from a broad candidate pool of 200 cytokines to a experimentally feasible data set, we subsequently applied an iterative PLSR filter method (Figure 3A) that reduced the full dataset from 200 cytokines to a 15 cytokine reduced model: CXCL15 (Accession #Q3UQ15), Marapsin (#Q14A25), DAN (#Q8C7N6), Fractalkine (#O35188), TGFβ-1 (#P04202), TROY (#Q80T13), IL-33 (#Q8BVZ5), Clusterin (#Q06890), MBL-2 (#P41317), Betacellulin (#Q05928), Chemerin (#Q8CHU8), CCL6 (#P27784), c-RP (#P14847), IL-7Ra (#Q9R0C1), MMP-3 (#P28862). While the full and reduced model were both 2-component models, the goodness-of-fit and predictive power (R^2^, Q^2^) increased in the reduced model (R^2^_reduced_ =0.607, Q^2^_reduced_ =0.555) compared to the full model (R^2^_full_ =0.166, Q^2^_full_ =0.414) (Figure 3B). The smaller discrepancy between R^2^ and Q^2^ and the increased Q^2^ also highlight the better predictive performance of the reduced model (69). The use of standard dimensionality reduction, PCA, and clustering on the full model dataset with or without preprocessing, does not show any biologically relevant clustering. However, analysis of the reduced dataset reveals clustering tightly connected to HSPC:MSPC seeding conditions, showing that the reduced model identified cytokine combinations that are distinguishable across culture conditions (Figure 4A,B).

**Figure 3.**
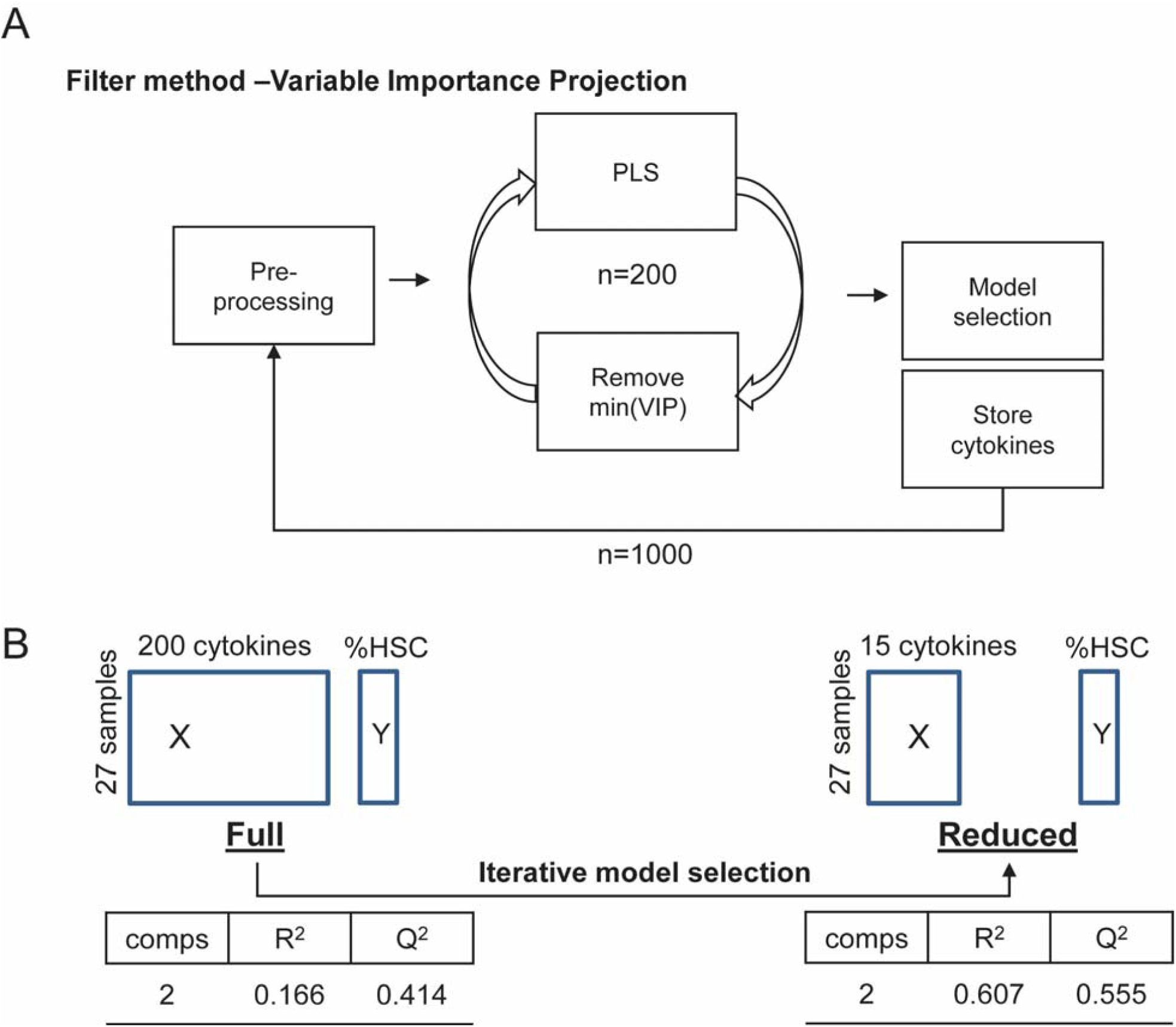
Partial Least Squares analysis of a scaled log-transformed dataset. **A.** The workflow of the iterative filter method to identify cytokines. The data is first scaled and log-transformed. Partial least squares regression (PLSR) is iteratively run on the preprocessed data, with the lowest variable importance projection (VIP) cytokine removed each time until all cytokines are removed (n=200). From this, the model with a Q^2^ within one S.E. of the maximum Q^2^ was chosen and the cytokines stored. This loop was then run 1,000 times. The cytokines that appear in >98% of the models were then chosen for the reduced model **B.** Model metrics of the full and reduced model are shown. The reduced model has a much smaller discrepancy between R^2^ and Q^2^.

**Figure 4.**
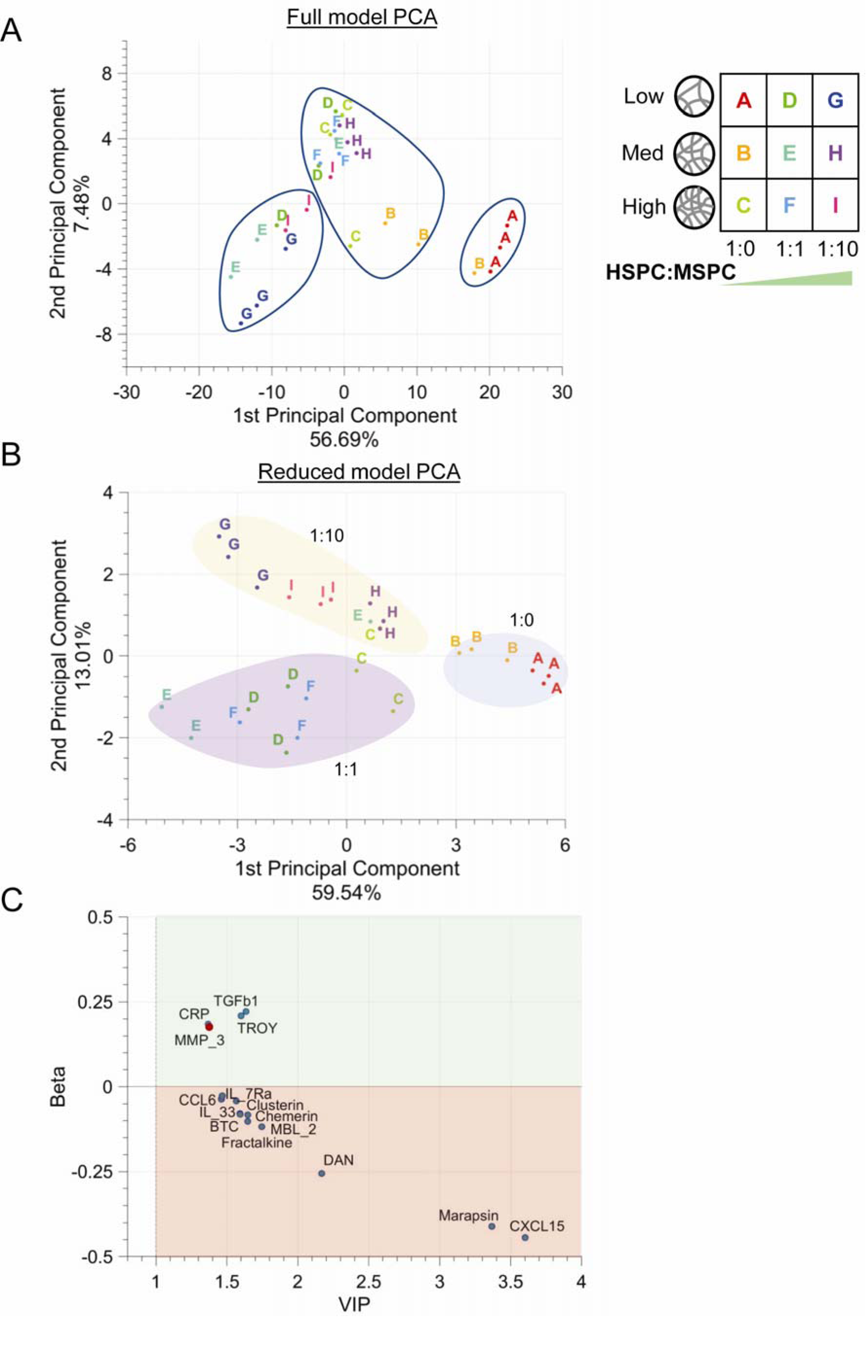
Model reduction and separation of regression coefficients. **A**. The scores of the first two components of principal component analysis (PCA) on the pre-processed data. Clustering is based on Euclidean distance and average linkage. The clusters do not capture either seeding or hydrogel conditions **B.** Scores of PCA on the pre-processed reduced data set. Clustering captures the seeding conditions with reasonable fidelity: Blue, 1:0, Purple 1:1, Yellow, 1:10 **C.** The scaled, centered, and log-transformed regression coefficients vs the variable importance projection. The upper quadrant (green) indicates a positive impact of the cytokines on HSC maintenance, while lower quadrant (red) indicates the reverse. MMP-3 data point is highlighted in red as it was significantly increased in the MSPC-only culture vs the HSPC-only culture, indicating it is a mainly paracrine signaling molecule.

### 3.3 Identifying secretome factors that correlate with hematopoietic cell response

We subsequently extracted transformed, scaled, and centered regression coefficients from the reduced cytokine model, plotting them against the variable importance projection (Figure 4C) (34). The positive or negative status of the regression coefficient split the reduced model cytokines into groups correlated with an increase (positive) or decrease (negative) in the HSC proportion. Given our goal to identify culture conditions for *ex vivo* expansion of hematopoietic progenitors, we subsequently concentrated on the family of cytokines correlated with a positive increase: TGFβ-1, MMP-3, TROY, and c-RP. However, future work may examine the role of cytokines negatively correlated with HSC maintenance as a means to potentially further manipulate HSC quiescence.

### 3.4 Validating secretome factor role via exogenous stimulation experiments

We subsequently performed new *in vitro* culture experiments to: 1) validate the effect of positively correlated cytokines on hematopoietic maintenance; and 2) to determine if the factors were likely acting directly on the HSPC population or indirectly through the MSPCs. To accomplish this, we exogenously stimulated both single HSPC cultures (1:0 HSPCs:MSPCs) and HSPC:MSPC co-cultures (1:1 HSPCs:MSPCs) encapsulated in 7.5 wt% GelMA hydrogels with individual positively correlated cytokines (TGFβ-1, MMP-3, c-RP, TROY) or a combination of all 4 cytokines. Latent MMP-3 and activated MMP-3 were tested separately as the microarray is unable to distinguish the two and have disparate functions. We examined the expansion of the combined population of long- and short-term hematopoietic stem cells (LSK, CD34^-/+^, CD135^-^) population. To represent the maintenance ability of the culture systems in response to exogenous stimulation, we have defined a metric, Progenitor Factor, as the ratio of early-stage progenitors to late-state committed progeny. The Progenitor Factor is the total (short-term plus long-term) HSC population normalized by the total population of lineage positive (differentiated) hematopoietic population and represents the ability of a culture to maintain a progenitor state (large Progenitor Factor) vs a committed state (small Progenitor Factor). The Progenitor Factor of each condition was normalized to the single-culture (HSPC-only), non-stimulated control. Regardless of the presence or absence of MSPCs, TGFβ-1, TROY, and the combination (TGFβ-1, MMP-3^latent^, TROY, and c-RP) increased the Progenitor Factor compared to the single-culture (HSPC only), non-stimulated control. The most notable increases in Progenitor Factor were in response to TGFβ-1, TROY, or Combination (with MMP-3^latent^). Single culture TGFβ-1, TROY, and the Combination (L) had a fold change of ΔPF_TGFβ-1_ = 8.73 ± 10.9, ΔPF_TROY_ = 9.33 ± 11.6, and ΔPF_Combo(L)_ = 7.52 ± 3.65 compared to the co-culture condition ΔPF_TGFβ-1_ = 6.07 ± 4.85, ΔPF_TROY_ = 0.874 ± 0.384, ΔPF_Combo(L)_ = 8.64 ± 4.31 (Figure 5A,B). The Glass’s Δ* effect size is a measure of the impact of a treatment compared to a control, which provides an estimate of the effect of each condition in order to identify conditions with the largest increase in hematopoietic potential from the non-stimulated, single-culture (Table 1).

**Figure 5.**
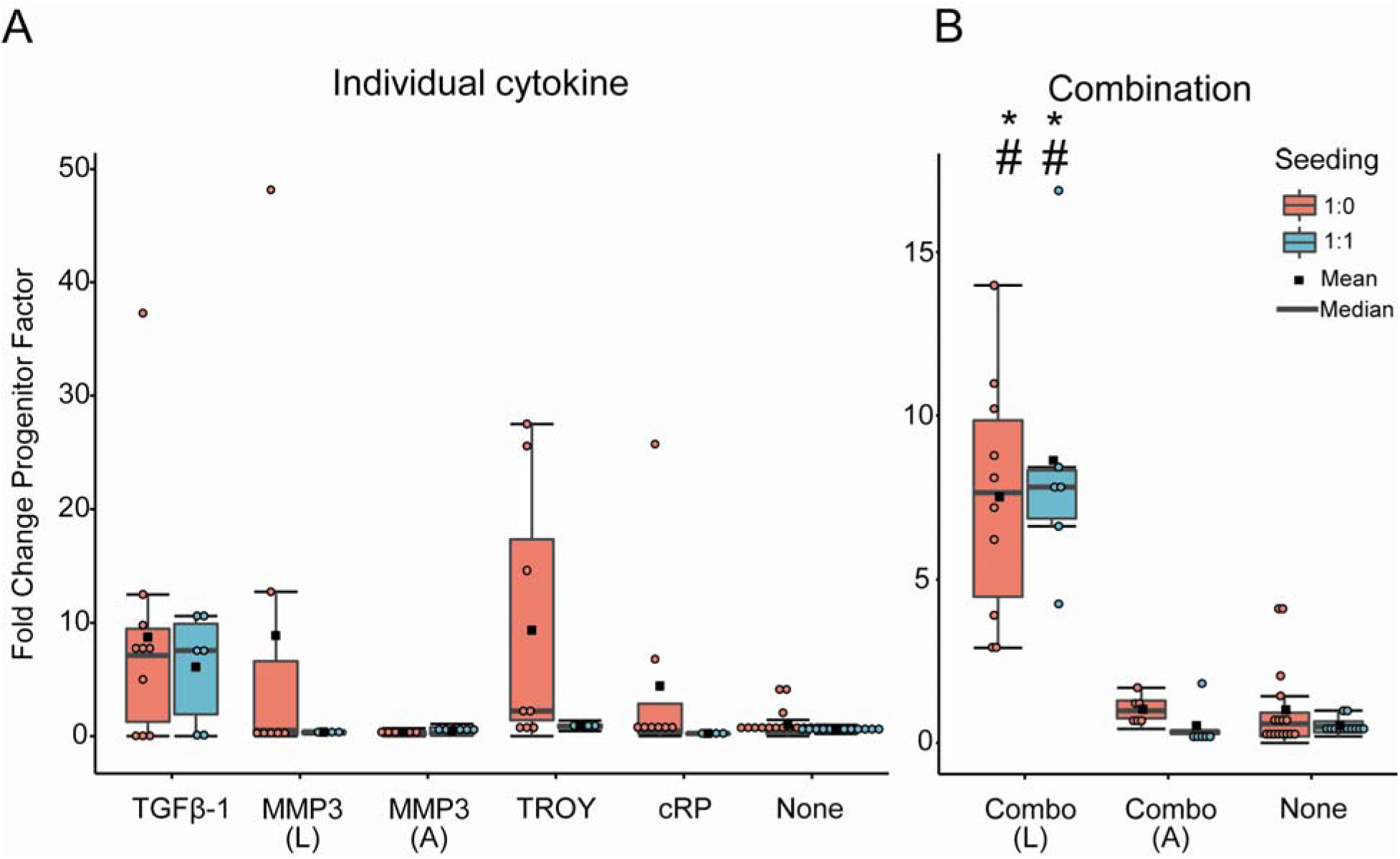
Hematopoietic maintenance potential in stimulated culture. Fold change of Progenitor Factor data. All conditions are normalized to the 1:0 None condition. # and * represent significance to 1:0 None and 1:1 None respectively (p<0.05). Each point represents one sample, with sample sizes listed below. (L) and (A) denote latent and activated MMP-3. Sample size is listed in Table 1. **A**. Progenitor Factor of individually stimulated conditions. **B.** Progenitor Factor of combination of cytokines.

### 3.5 The role of exogenous stimulation HSPC quiescence

Finally, we examined the influence of model-identified cytokine stimulation on the maintenance of quiescent HSPCs within the hydrogel culture. Quiescent HSCs are essential in maintaining the hematopoietic population during homeostasis and injury. The number of quiescent (G0) HSCs was dependent upon both the seeding condition and cytokine stimulus and is reported as a fold change from the non-stimulated HSPC-only culture (Figure 6). Exogenous addition of Combination^latent^ (TGFβ-1, latent MMP3, TROY, cRP) led to a significant (p-value < 0.05) fold increase in the number of quiescent cells (ΔG0_Combo(L)_ 5.59 ± 6.17 for HSPC only, 6.02 ± 1.32 for HSPC:MSPC co-culture) in comparison to the unstimulated controls for both single-(HSPC only) and co-cultures (1:1 HSPC:MSPC). This increase is also observed with exogenous addition of TGFβ-1 only (ΔG0_TGFβ-1_: 5.0 ± 7.36 for HSPC-only single cultures; 5.70 ± 4.14 for 1:1 HSPC:MSPC co-cultures), albeit with greater variability. Notably, the presence of MSPCs is important for observed increase in ΔG0 cells, as all exogenous factor conditions experienced a higher ΔG0 with the presence of MPSCs (Figure 6). We also see that the presence of MSPCs leads to a greater proportion of HSCs that are in the G0 phase (Supp. Figure 2). Taken together, this shows that while the overall shift in the hematopoietic population (Progenitor Factor) is independent of the presence or absence of MSPCs, the number of quiescent hematopoietic cells is increased by co-culture with MSPCs stimulated with exogenous factors (Figure 5,6).

**Figure 6.**
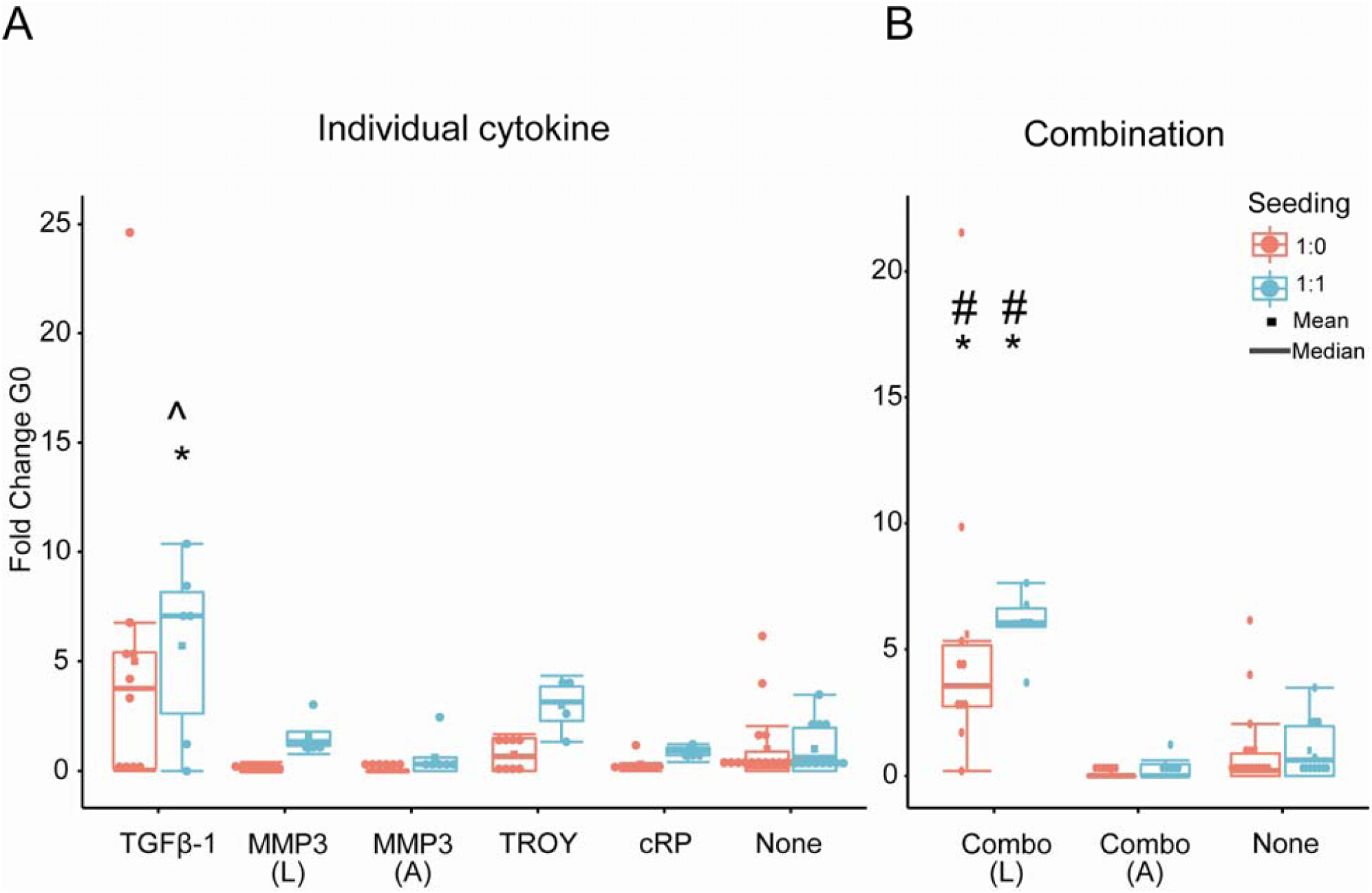
Quiescent HSCs in stimulated cultured. Fold change of quiescent (G0) HSCs, normalized to the 1:0 None condition. # and * represent significance to 1:0 None and 1:1 None respectively (p<0.05), ^ represents significance to 1:1 None at p<0.1 (p=0.079). Each point represents one sample, with sample sizes listed below. (L) and (A) denote latent and activated MMP-3. **A.** Fold change of quiescent HSCs in individually stimulated conditions. **B.** Fold change of quiescent HSCs exposed to combination of cytokines.

## 4. Discussion

The hematopoietic stem cell microenvironment in the bone marrow has inspired the concept of the stem cell niche (70) as well as experimental efforts to identify the role of biophysical and biomolecular elements in the niche. One major axis of investigation regarding signals within the HSC niche are the multitude of signaling biomolecules originating from the many cell lineages that exist within the marrow, including cells of the hematopoietic and mesenchymal progeny (13, 14). While the exact source of soluble factors in the native bone marrow environment can be troublesome to elucidate, the cell-secreted factors are implicated in *in vivo* and *ex vivo* maintenance of hematopoietic activity (71). Co-transplantation of HSC and mesenchymal stromal cells (MSCs) improves hematopoietic engraftment and feeder-layers of mesenchymal progeny leads to *ex vivo* maintenance. However, the native complexity and imaging challenges of these studies hinders *in vivo* elucidation of influential HSC-MSC interactions. Previously, our lab has shown that the presence of MSPCs in GelMA hydrogels increases hematopoietic maintenance of HSPCs compared to single-cultures. The presence of MSPCs led to long-range cell-cell communication via secretion of biomolecules related to cell signaling (cytokines) and non-uniform remodeling of the local and non-local matrix (matrix metalloproteases). To quantify the influence of cell-secreted factors on HSC maintenance, we examined the secretome of media conditioned by HSPCs and MSPCs in single or co-cultures via a quantitative cytokine microarray. While conditioned media does not allow for quantification of soluble factors that have been sequestered within the GelMA hydrogel, it does offer an accessible avenue for probing the secretome.

Analysis of the secretome yielded a large and complex dataset that necessitated reduction and visualization techniques to parse out relevant trends within the data. While traditional dimensional analysis techniques (heatmap, PCA) can reveal clustering and differential concentrations (Figure 2), they lack the ability to correlate measured observations with a biological process. These methods are also sensitive to noise within the data and can lead to results that do not have a biological significance. As such, another analysis technique was required to shift through the large dataset to identify cytokines that are relevant to maintaining HSCs within an engineered niche. For this, we used a modeling technique that has been employed in analysis of signaling pathways and metabolomics: Partial Least Squares Regression (PLSR) (72-74). Similar to PCA, PLSR reduces the dimensionality of complex data; however, PLSR has the advantage of correlating observational data (secretome) to a biological process (HSC maintenance) and is better able to distinguish signal from noise (56, 59, 75-77). But while PLSR is able to correlate concentration of soluble factors to hematopoietic phenotype, an additional variable selection step is required to minimize the number of cytokines implicated in HSC maintenance. We used an iterative filter method, where the cytokine with the lowest variable importance projection (VIP; a measure of the importance of the cytokine to the model) was removed from the model. VIP was selected as the filter criteria as it is considered a stable filter method and produces consistent results (78). The repeated use (n=1,000) of this iterative PLSR filter method for variable selection resulted in a reduced candidate pool of 15 cytokines that were correlated with HSC maintenance. PLSR has previously been applied to stem cell systems, including exploration of soluble factor signaling, with excellent work by Müller et al. applying PLSR to a hematopoietic system to implicate autocrine, paracrine, and juxtacrine signals in HSPC proliferation (34, 79, 80). While Müller et al. used a cutoff of VIP>0.8 in their final model to create an optimized, reduced model, it is not apparent that variable selection methods, beyond final cutoff values, have been used to identify minimum cocktails of cytokines that impact stem cell response. Although variable selection methods have been employed in chemometric analysis to produce models with better overall predictive ability (R^2^ and Q^2^) (81, 82), our effort represents an important alignment of PLSR methods with stem cell culture platforms to identify essential cell-cell interactions that underlie stem cell performance.

Our approach also highlights the synergy between bioinformatics and experimental approaches, in which we subsequently examined the individual and combined effects of cytokines positively correlated to improved HSPC response (TGFβ-1, MMP-3, c-RP, TROY) in mixed HSPC-MSPC cultures. One of the challenges associated with experimental design of exogenous stimulation of stem cells is identifying cytokine dosages that trigger a biological response. While the measured concentration of cytokines ranges from 1 to 15 ng/mL, we used a consistent concentration of 50 ng/mL for all factors in order to identify biological effects. The higher dose does not mitigate inherent variability in *in vitro* culture and biological response, and future studies may explore dose dependent responses amongst these factors. However, this approach does demonstrate the utility of a robust protocol to identify statistically significant stem cell response. As PLSR is unable to distinguish the source of the identified cytokines, it was not known whether the cytokines act directly on the hematopoietic population or indirectly via MSPC activation. We performed cytokine stimulation experiments on both single (HSPC) and HSPC-MSPC co-cultures, finding that improvements in HSPC Progenitor Factor was independent of the MSPC seeding condition, i.e. conditions have similar calculated Progenitor Factors regardless of the presence of MSPCs (Figure 5). This suggests that the exogenously added cytokines are likely acting directly upon the HSCs. The highest Progenitor Factor was observed with the combination of all 4 factors (latent MMP-3), with a large effect size (>30) compared to the non-stimulated control, suggesting a possible interaction effect among the cytokines (Figure 5B, Table 1). This is supported by STRING protein-protein association analysis which shows a complex network of co-expression and known interactions (83), and opens the door to subsequent use of our HSPC culture platform to more explicitly probe such interactions (Supp. Figure 3).

While the Progenitor Factor appears independent of the presence of MSPCs, the HSC quiescent state is sensitive to both exogenous stimulation and seeding condition. Exogenous stimulation with TGFβ-1 (alone or in combination) led to a 5-fold increase in the number of quiescent HSCs in HSPC-only (single) and 1:1 HSPC-MSPC (co-culture) cultures (Figure 6). Excitingly, this *in vitro* response is reminiscent of proposed features of the *in vivo* niche, where quiescent HSCs are often found near megakaryocytes, a major producer of TGFβ-1 (14). Further, the presence of MSPCs led to a higher number of G0 cells in all conditions, compared to single-cultures. Taken together, this suggests that lineage patterns are independent of seeding density when stimulated with a combination of exogenous cytokines, yet HSC cell-cycle state appears to be dominated by MSPC-mediated interactions. This highlights both the complexity of a synthetic HSC niche, and also the potential for experimental-modeling approaches to identify and manipulate heterotypic cell interactions in order to regulate a desired stem cell response. Our approach provides critical information about the potential for HSPC-MSPC interactions to achieve expansion of an HSPC population without exhaustion (e.g., maintenance of a quiescent cell fraction).

We have noted that TGFβ-1 response *in vitro* can be linked to an *in vivo* megakaryocyte niche, and this is mirrored by c-RP which has been suggested to stimulate the production of megakaryocytes (megakaryocytopoiesis) (84). The exact functionality of c-RP *in vivo* and *in vitro* remains elusive (85), however, it has been reported as a key regulator of systemic inflammation and is a prognosis biomarker for HSC transplant success (86, 87). Similarly, TROY (tumor necrosis factor receptor super family 19) has not been fully explored in the context of hematopoiesis, although other members of the TNFR family have been related to the emergence of hematopoietic stem cells and immunomodulatory roles (88). Interestingly, it has been shown to regulate MSC differentiation (89), indicating a route by which TROY can indirectly impact HSC maintenance. However, our approach is unique in that it took an experimental co-culture, a broad secretome screen, then an adaptation of conventional PLSR approached to identify select factors involved in HSPC expansion and quiescence in a synthetic HSC niche. It is exciting that many of these factors identified via this approach have known hematopoietic or bone marrow origin, suggesting this approach may have broad value for refining an *ex vivo* platform for HSC culture and expansion.

Importantly, while soluble factors can act directly upon cultured HSPCs and MSPCs, secretome analysis also enables examination of the effect of dynamic changes to the matrix by MSPCs. MSPCs can participate in matrix remodeling via the secretion of degradative enzymes (MMPs) and their inhibitors (TIMPs). In the reduced model, MMP-3 is positively correlated and is primarily a paracrine signaling molecule (Figure 4C), with significantly higher concentrations in the MSPC single cultures compared to the HSPC single cultures. MMP-3 degrades a wide variety of ECM components including collagen type II, IV, V, IX (90) and serves as an activator for other MMPs, setting off a remodeling cascade (91-94). While this model has not explored the activity of MMP-3 or its inhibitors, it does suggest that there is a need for deeper examination of MMPs and TIMPs and their associated role in matrix remodeling.

This work has validated the influence of soluble factors in maintaining a progenitor population in an *in vitro* niche. With PLSR, we have shown that a complex secretome dataset can be correlated to a biological response for hematopoietic stem cell engineering. A minimum cocktail of cytokines identified by the model was shown to increase the progenitor potential of the culture system, acting directly upon the hematopoietic population to maintain a higher Progenitor Factor. As this work was performed at cytokine concentrations of 50 ng/mL, there is the possibility that the high concentration of each cytokine hid any indirect effects arising from MSPC-secreted factors. However, reducing the pool of candidate cytokines from 200 to 4 now enables future dose-dependent and temporal secretion studies of HSC lineage patterns. Additionally, this small subset allows for future inhibition studies that further solidify the role of these factors in hematopoietic maintenance or that examine negatively correlated cytokines (e.g. Marapsin). In parallel, future work will probe changes in long-range cell-cell communication mediated by changes in the local and non-local matrix. Following the method outlined herein, remodeling-associated factors within the secretome can be similarly correlated to bulk-material mechanical measurements (compression testing) and local matrix remodeling (microrheology (95)). The sum total of this work will identify factors implicated in the maintenance of a hematopoietic stem cell population and inform design and development of an artificial stem cell niche.

## Acknowledgments

The authors would like to acknowledge Dr. Barbara Pilas and Barbara Balhan of the Roy J. Carver Biotechnology Center (Flow Cytometry Facility, UIUC) for assistance with bone marrow cell isolation and flow cytometry. Research reported in this publication was supported by the National Institute of Diabetes and Digestive and Kidney Diseases of the National Institutes of Health under Award Numbers R01 DK099528 (B.A.C.H) and F31 DK117514 (A.E.G.), as well as by the National Institute of Biomedical Imaging and Bioengineering of the National Institutes of Health under Award Numbers R21 EB018481 (B.A.C.H.) and T32 EB019944 (A.E.G.). The content is solely the responsibility of the authors and does not necessarily represent the official views of the NIH. The authors are also grateful for additional funding provided by the Department of Chemical & Biomolecular Engineering and the Institute for Genomic Biology at the University of Illinois at Urbana-Champaign.

## References

1. Yang L, Bryder D, Adolfsson J, Nygren J, Mansson R, Sigvardsson M, et al. Identification of Lin(-)Sca1(+)kit(+)CD34(+)Flt3-short-term hematopoietic stem cells capable of rapidly reconstituting and rescuing myeloablated transplant recipients. Blood. 2005;105(7):2717–23.

2. Adams GB, Scadden DT. The hematopoietic stem cell in its place. Nat Immunol. 2006;7(4):333–7.

3. Boulais PE, Frenette PS. Making sense of hematopoietic stem cell niches. Blood. 2015;125(17):2621–9.

4. Calvi LM, Adams GB, Weibrecht KW, Weber JM, Olson DP, Knight MC, et al. Osteoblastic cells regulate the haematopoietic stem cell niche. Nature. 2003;425(6960):841–6.

5. Hines M, Nielsen L, Cooper-White J. The hematopoietic stem cell niche: what are we trying to replicate? Journal of Chemical Technology & Biotechnology. 2008;83(4):421–43.

6. Krause DS, Scadden DT, Preffer FI. The hematopoietic stem cell niche--home for friend and foe? Cytometry B Clin Cytom. 2013;84(1):7–20.

7. Mendez-Ferrer S, Lucas D, Battista M, Frenette PS. Haematopoietic stem cell release is regulated by circadian oscillations. Nature. 2008;452(7186):442–7.

8. Morrison SJ, Scadden DT. The bone marrow niche for haematopoietic stem cells. Nature. 2014;505(7483):327–34.

9. Pinho S, Marchand T, Yang E, Wei Q, Nerlov C, Frenette PS. Lineage-Biased Hematopoietic Stem Cells Are Regulated by Distinct Niches. Dev Cell. 2018;44(5):634–41 e4.

10. Purton LE, Scadden DT. The hematopoietic stem cell niche. StemBook. Cambridge (MA) 2008.

11. Zhang J, Li L. Stem cell niche: microenvironment and beyond. J Biol Chem. 2008;283(15):9499–503.

12. Jansen LE, Birch NP, Schiffman JD, Crosby AJ, Peyton SR. Mechanics of intact bone marrow. J Mech Behav Biomed Mater. 2015;50:299–307.

13. Wei Q, Frenette PS. Niches for Hematopoietic Stem Cells and Their Progeny. Immunity. 2018;48(4):632–48.

14. Pinho S, Frenette PS. Haematopoietic stem cell activity and interactions with the niche. Nat Rev Mol Cell Biol. 2019;20(5):303–20.

15. Mendelson A, Frenette PS. Hematopoietic stem cell niche maintenance during homeostasis and regeneration. Nat Med. 2014;20(8):833–46.

16. Nombela-Arrieta C, Silberstein LE. The science behind the hypoxic niche of hematopoietic stem and progenitors. Hematology Am Soc Hematol Educ Program. 2014;2014(1):542–7.

17. Spencer JA, Ferraro F, Roussakis E, Klein A, Wu J, Runnels JM, et al. Direct measurement of local oxygen concentration in the bone marrow of live animals. Nature. 2014;508(7495):269–73.

18. Ganuza M, McKinney-Freeman S. Hematopoietic stem cells under pressure. Curr Opin Hematol. 2017;24(4):314–21.

19. Shono Y, Ueha S, Wang Y, Abe J, Kurachi M, Matsuno Y, et al. Bone marrow graft-versus-host disease: early destruction of hematopoietic niche after MHC-mismatched hematopoietic stem cell transplantation. Blood. 2010;115(26):5401–11.

20. National Institutes of Health USDoHaHS. NIH Stem Cell Information Home Page 2016 [Available from: stemcells.nih.gov/info/2001report/chapter5.htm.

21. Wilkinson AC, Ishida R, Kikuchi M, Sudo K, Morita M, Crisostomo RV, et al. Long-term ex vivo haematopoietic-stem-cell expansion allows nonconditioned transplantation. Nature. 2019;571(7763):117–21.

22. Wolff SN. Second hematopoietic stem cell transplantation for the treatment of graft failure, graft rejection or relapse after allogeneic transplantation. Bone Marrow Transplant. 2002;29(7):545–52.

23. Wekerle T, Kurtz J, Ito H, Ronquillo JV, Dong V, Zhao G, et al. Allogeneic bone marrow transplantation with co-stimulatory blockade induces macrochimerism and tolerance without cytoreductive host treatment. Nat Med. 2000;6(4):464–9.

24. Donnelly H, Salmeron-Sanchez M, Dalby MJ. Designing stem cell niches for differentiation and self-renewal. J R Soc Interface. 2018;15(145).

25. Zhang Z, Gupte MJ, Ma PX. Biomaterials and stem cells for tissue engineering. Expert Opin Biol Ther. 2013;13(4):527–40.

26. Lutolf MP, Gilbert PM, Blau HM. Designing materials to direct stem-cell fate. Nature. 2009;462(7272):433–41.

27. Choi JS, Mahadik BP, Harley BA. Engineering the hematopoietic stem cell niche: Frontiers in biomaterial science. Biotechnol J. 2015;10(10):1529–45.

28. Lee-Thedieck C, Rauch N, Fiammengo R, Klein G, Spatz JP. Impact of substrate elasticity on human hematopoietic stem and progenitor cell adhesion and motility. J Cell Sci. 2012;125(Pt 16):3765–75.

29. Chaudhuri O, Gu L, Klumpers D, Darnell M, Bencherif SA, Weaver JC, et al. Hydrogels with tunable stress relaxation regulate stem cell fate and activity. Nat Mater. 2016;15(3):326–34.

30. Deans TL, Singh A, Gibson M, Elisseeff JH. Regulating synthetic gene networks in 3D materials. Proc Natl Acad Sci U S A. 2012;109(38):15217–22.

31. Singh A, Deans TL, Elisseeff JH. Photomodulation of Cellular Gene Expression in Hydrogels. Acs Macro Letters. 2013;2(3):269–72.

32. Choi JS, Harley BA. Marrow-inspired matrix cues rapidly affect early fate decisions of hematopoietic stem and progenitor cells. Sci Adv. 2017;3(1):e1600455.

33. Mahadik BP, Pedron Haba S, Skertich LJ, Harley BA. The use of covalently immobilized stem cell factor to selectively affect hematopoietic stem cell activity within a gelatin hydrogel. Biomaterials. 2015;67:297–307.

34. Muller E, Wang W, Qiao W, Bornhauser M, Zandstra PW, Werner C, et al. Distinguishing autocrine and paracrine signals in hematopoietic stem cell culture using a biofunctional microcavity platform. Sci Rep. 2016;6:31951.

35. Di Maggio N, Piccinini E, Jaworski M, Trumpp A, Wendt DJ, Martin I. Toward modeling the bone marrow niche using scaffold-based 3D culture systems. Biomaterials. 2011;32(2):321–9.

36. Csaszar E, Kirouac DC, Yu M, Wang W, Qiao W, Cooke MP, et al. Rapid expansion of human hematopoietic stem cells by automated control of inhibitory feedback signaling. Cell Stem Cell. 2012;10(2):218–29.

37. Cambier T, Honegger T, Vanneaux V, Berthier J, Peyrade D, Blanchoin L, et al. Design of a 2D no-flow chamber to monitor hematopoietic stem cells. Lab Chip. 2015;15(1):77–85.

38. Stylianopoulos T, Poh MZ, Insin N, Bawendi MG, Fukumura D, Munn LL, et al. Diffusion of particles in the extracellular matrix: the effect of repulsive electrostatic interactions. Biophys J. 2010;99(5):1342–9.

39. Kamali-Zare P, Nicholson C. Brain extracellular space: geometry, matrix and physiological importance. Basic Clin Neurosci. 2013;4(4):282–6.

40. Pluen A, Netti PA, Jain RK, Berk DA. Diffusion of macromolecules in agarose gels: comparison of linear and globular configurations. Biophys J. 1999;77(1):542–52.

41. van Donkelaar CC, Chao G, Bader DL, Oomens CW. A reaction-diffusion model to predict the influence of neo-matrix on the subsequent development of tissue-engineered cartilage. Comput Methods Biomech Biomed Engin. 2011;14(5):425–32.

42. Sakiyama-Elbert SE. Incorporation of heparin into biomaterials. Acta Biomater. 2014;10(4):1581–7.

43. Mahadik BP, Bharadwaj NA, Ewoldt RH, Harley BA. Regulating dynamic signaling between hematopoietic stem cells and niche cells via a hydrogel matrix. Biomaterials. 2017;125:54–64.

44. Leisten I, Kramann R, Ventura Ferreira MS, Bovi M, Neuss S, Ziegler P, et al. 3D co-culture of hematopoietic stem and progenitor cells and mesenchymal stem cells in collagen scaffolds as a model of the hematopoietic niche. Biomaterials. 2012;33(6):1736–47.

45. Gvaramia D, Muller E, Muller K, Atallah P, Tsurkan M, Freudenberg U, et al. Combined influence of biophysical and biochemical cues on maintenance and proliferation of hematopoietic stem cells. Biomaterials. 2017;138:108–17.

46. Gilchrist AE, Lee S, Hu Y, Harley BAC. Soluble Signals and Remodeling in a Synthetic Gelatin-Based Hematopoietic Stem Cell Niche. Adv Healthc Mater. 2019;8(20):e1900751.

47. Okada S, Nakauchi H, Nagayoshi K, Nishikawa SI, Miura Y, Suda T. Invivo and Invitro Stem-Cell Function of C-Kit-Positive and Sca-1-Positive Murine Hematopoietic-Cells. Blood. 1992;80(12):3044–50.

48. Challen GA, Boles N, Lin KK, Goodell MA. Mouse hematopoietic stem cell identification and analysis. Cytometry A. 2009;75(1):14–24.

49. Benton JA, DeForest CA, Vivekanandan V, Anseth KS. Photocrosslinking of gelatin macromers to synthesize porous hydrogels that promote valvular interstitial cell function. Tissue Eng Part A. 2009;15(11):3221–30.

50. Pedron S, Harley BA. Impact of the biophysical features of a 3D gelatin microenvironment on glioblastoma malignancy. J Biomed Mater Res A. 2013;101(12):3404–15.

51. Chen JE, Pedron S, Harley BAC. The Combined Influence of Hydrogel Stiffness and Matrix-Bound Hyaluronic Acid Content on Glioblastoma Invasion. Macromol Biosci. 2017;17(8).

52. Feng C, Wang H, Lu N, Chen T, He H, Lu Y, et al. Log-transformation and its implications for data analysis. Shanghai Arch Psychiatry. 2014;26(2):105–9.

53. Ekwaru JP, Veugelers PJ. The Overlooked Importance of Constants Added in Log Transformation of Independent Variables with Zero Values: A Proposed Approach for Determining an Optimal Constant. Statistics in Biopharmaceutical Research. 2017;10(1):26–9.

54. AB SSDA. User Guide SIMCA® 15 Multivariate Data Analysis Solution 2017.

55. van den Berg RA, Hoefsloot HC, Westerhuis JA, Smilde AK, van der Werf MJ. Centering, scaling, and transformations: improving the biological information content of metabolomics data. BMC Genomics. 2006;7:142.

56. Worley B, Powers R. Multivariate Analysis in Metabolomics. Curr Metabolomics. 2013;1(1):92–107.

57. Team R. RStudio: Integrated Development Environment for R. RStudio, Inc.; 2019.

58. Rohart F, Gautier B, Singh A, Le Cao KA. mixOmics: An R package for ‘omics feature selection and multiple data integration. PLoS Comput Biol. 2017;13(11):e1005752.

59. Jorgensen H, Hill AS, Beste MT, Kumar MP, Chiswick E, Fedorcsak P, et al. Peritoneal fluid cytokines related to endometriosis in patients evaluated for infertility. Fertil Steril. 2017;107(5):1191–9 e2.

60. D’Haeseleer P. How does gene expression clustering work? Nat Biotechnol. 2005;23(12):1499–501.

61. Yang LP, Bryder D, Adolfsson J, Nygren J, Mansson R, Sigvardsson M, et al. Identification of Lin(-)Sca1(+)kit(+)CD34(+)Flt(3-) short-term hematopoietic stem cells capable of rapidly reconstituting and rescuing myeloablated transplant recipients. Blood. 2005;105(7):2717–23.

62. Beaudin AE, Boyer SW, Forsberg EC. Flk2/Flt3 promotes both myeloid and lymphoid development by expanding non-self-renewing multipotent hematopoietic progenitor cells. Exp Hematol. 2014;42(3):218–29 e4.

63. Tian C, Zhang Y. Purification of hematopoietic stem cells from bone marrow. Ann Hematol. 2016;95(4):543–7.

64. Zhang Y, Yan X, Sashida G, Zhao X, Rao Y, Goyama S, et al. Stress hematopoiesis reveals abnormal control of self-renewal, lineage bias, and myeloid differentiation in Mll partial tandem duplication (Mll-PTD) hematopoietic stem/progenitor cells. Blood. 2012;120(5):1118–29.

65. Brown MB, Forsythe AB. Robust Tests for the Equality of Variances. Journal of the American Statistical Association. 1974;69(346):364–7.

66. Ghasemi A, Zahediasl S. Normality tests for statistical analysis: a guide for non-statisticians. Int J Endocrinol Metab. 2012;10(2):486–9.

67. Shingala MC, Rajyaguru A. Comparison of Post Hoc Tests for Unequal Variance. International Journal of New Technologies in Science and Engineering. 2015;2(5).

68. Ialongo C. Understanding the effect size and its measures. Biochem Med (Zagreb). 2016;26(2):150–63.

69. Triba MN, Le Moyec L, Amathieu R, Goossens C, Bouchemal N, Nahon P, et al. PLS/OPLS models in metabolomics: the impact of permutation of dataset rows on the K-fold cross-validation quality parameters. Mol Biosyst. 2015;11(1):13–9.

70. Schofield R. The relationship between the spleen colony-forming cell and the haemopoietic stem cell. Blood cells. 1978;4(1-2):7–25.

71. Sorrentino BP. Clinical strategies for expansion of haematopoietic stem cells. Nat Rev Immunol. 2004;4(11):878–88.

72. Wold S, Sjostrom M, Eriksson L. PLS-regression: a basic tool of chemometrics. Chemometrics and Intelligent Laboratory Systems. 2001;58(2):109–30.

73. Janes KA, Kelly JR, Gaudet S, Albeck JG, Sorger PK, Lauffenburger DA. Cue-signal-response analysis of TNF-induced apoptosis by partial least squares regression of dynamic multivariate data. J Comput Biol. 2004;11(4):544–61.

74. Kinney MA, Vo LT, Frame JM, Barragan J, Conway AJ, Li S, et al. A systems biology pipeline identifies regulatory networks for stem cell engineering. Nat Biotechnol. 2019;37(7):810–8.

75. Kreeger PK. Using partial least squares regression to analyze cellular response data. Sci Signal. 2013;6(271):tr7.

76. Plantier L, Cazes A, Dinh-Xuan AT, Bancal C, Marchand-Adam S, Crestani B. Physiology of the lung in idiopathic pulmonary fibrosis. Eur Respir Rev. 2018;27(147).

77. Bartel J, Krumsiek J, Theis FJ. Statistical methods for the analysis of high-throughput metabolomics data. Comput Struct Biotechnol J. 2013;4:e201301009.

78. Mehmood T, Liland KH, Snipen L, Saebo S. A review of variable selection methods in Partial Least Squares Regression. Chemometrics and Intelligent Laboratory Systems. 2012;118:62–9.

79. Chai YC, Roberts SJ, Desmet E, Kerckhofs G, van Gastel N, Geris L, et al. Mechanisms of ectopic bone formation by human osteoprogenitor cells on CaP biomaterial carriers. Biomaterials. 2012;33(11):3127–42.

80. Peltier J, Schaffer DV. Systems biology approaches to understanding stem cell fate choice. IET Syst Biol. 2010;4(1):1–11.

81. Goodarzi M, Dejaegher B, Vander Heyden Y. Feature selection methods in QSAR studies. J AOAC Int. 2012;95(3):636–51.

82. Chong IG, Jun CH. Performance of some variable selection methods when multicollinearity is present. Chemometrics and Intelligent Laboratory Systems. 2005;78(1-2):103–12.

83. Szklarczyk D, Gable AL, Lyon D, Junge A, Wyder S, Huerta-Cepas J, et al. STRING v11: protein-protein association networks with increased coverage, supporting functional discovery in genome-wide experimental datasets. Nucleic Acids Res. 2019;47(D1):D607–D13.

84. Potempa LA, Motie M, Wright KE, Crump BL, Radosevich JA, Sakai N, et al. Stimulation of megakaryocytopoiesis in mice by human modified C-reactive protein (mCRP). Exp Hematol. 1996;24(2):258–64.

85. Potempa LA, Yao ZY, Ji SR, Filep JG, Wu Y. Solubilization and purification of recombinant modified C-reactive protein from inclusion bodies using reversible anhydride modification. Biophys Rep. 2015;1:18–33.

86. Min CK, Kim SY, Eom KS, Kim YJ, Kim HJ, Lee S, et al. Patterns of C-reactive protein release following allogeneic stem cell transplantation are correlated with leukemic relapse. Bone Marrow Transplant. 2006;37(5):493–8.

87. Jordan KK, Christensen IJ, Heilmann C, Sengelov H, Muller KG. Pretransplant C-reactive protein as A prognostic marker in allogeneic stem cell transplantation. Scand J Immunol. 2014;79(3):206–13.

88. Denk A, Wirth T, Baumann B. NF-κB transcription factors: critical regulators of hematopoiesis and neuronal survival. Cytokine & Growth Factor Reviews. 2000;11(4):303–20.

89. Qiu W, Hu Y, Andersen TE, Jafari A, Li N, Chen W, et al. Tumor necrosis factor receptor superfamily member 19 (TNFRSF19) regulates differentiation fate of human mesenchymal (stromal) stem cells through canonical Wnt signaling and C/EBP. J Biol Chem. 2010;285(19):14438–49.

90. Ennis BW, Matrisian LM. Matrix degrading metalloproteinases. J Neurooncol. 1994;18(2):105–9.

91. Johnson JL, Dwivedi A, Somerville M, George SJ, Newby AC. Matrix metalloproteinase (MMP)-3 activates MMP-9 mediated vascular smooth muscle cell migration and neointima formation in mice. Arterioscler Thromb Vasc Biol. 2011;31(9):e35–44.

92. Ye S, Eriksson P, Hamsten A, Kurkinen M, Humphries SE, Henney AM. Progression of coronary atherosclerosis is associated with a common genetic variant of the human stromelysin-1 promoter which results in reduced gene expression. J Biol Chem. 1996;271(22):13055–60.

93. Toth M, Chvyrkova I, Bernardo MM, Hernandez-Barrantes S, Fridman R. Pro-MMP-9 activation by the MT1-MMP/MMP-2 axis and MMP-3: role of TIMP-2 and plasma membranes. Biochem Biophys Res Commun. 2003;308(2):386–95.

94. Dreier R, Grassel S, Fuchs S, Schaumburger J, Bruckner P. Pro-MMP-9 is a specific macrophage product and is activated by osteoarthritic chondrocytes via MMP-3 or a MT1-MMP/MMP-13 cascade. Exp Cell Res. 2004;297(2):303–12.

95. Schultz KM, Kyburz KA, Anseth KS. Measuring dynamic cell-material interactions and remodeling during 3D human mesenchymal stem cell migration in hydrogels. Proc Natl Acad Sci U S A. 2015;112(29):E3757–64.

