## supplemental for "Connecting secretome to hematopoietic stem cell phenotype shifts in an engineered bone marrow niche"

**Supplemental Table 1.** Complete list of 200 cytokines included in the RayBiotech microarray panel ((#QAM-CAA-4000-1) including common protein names, gene symbol, UniProt Accession#, and UniProt reference link.

| **Protein Name** | **Gene Symbol** | **UniProt Accession #** | **UniProt URL** |
| --- | --- | --- | --- |
| 4-1BB | TNFRSF9 | P20334 | http://www.uniprot.org/uniprot/P20334 |
| 6Ckine | CCL21 | P86792 | http://www.uniprot.org/uniprot/P86792 |
| Ace | ACE | P22967 | http://www.uniprot.org/uniprot/P22967 |
| Activin A | INHBA | Q04998 | http://www.uniprot.org/uniprot/Q04998 |
| ADAMTS-1 | ADAMTS1 | P97857 | http://www.uniprot.org/uniprot/P97857 |
| Adiponectin | ADIPOQ | Q62400 | http://www.uniprot.org/uniprot/Q62400 |
| ALK-1 | SLPI | O09081 | http://www.uniprot.org/uniprot/O09081 |
| Amphiregulin | AREG | P31955 | http://www.uniprot.org/uniprot/P31955 |
| ANGPTL1 | ANGPTL1 | Q640P2 | http://www.uniprot.org/uniprot/Q640P2 |
| ANGPTL3 | ANGPTL3 | Q9R182 | http://www.uniprot.org/uniprot/Q9R182 |
| Artemin | ARTN | Q3SXF4 | http://www.uniprot.org/uniprot/Q3SXF4 |
| Axl | Axl | Q00993 | http://www.uniprot.org/uniprot/Q00993 |
| BAFF R | TNFRSF13C | Q9D8D0 | http://www.uniprot.org/uniprot/Q9D8D0 |
| Betacellulin | BTC | Q05928 | http://www.uniprot.org/uniprot/Q05928 |
| bFGF | FGF2 | P15655 | http://www.uniprot.org/uniprot/P15655 |
| BLC | CXCL13 | O55038 | http://www.uniprot.org/uniprot/O55038 |
| C5a | C5 | P06684 | http://www.uniprot.org/uniprot/P06684 |
| Cardiotrophin-1 | CTF1 | Q60753 | http://www.uniprot.org/uniprot/Q60753 |
| CCL28 | CCL28 | Q9JIL2 | http://www.uniprot.org/uniprot/Q9JIL2 |
| CCL6 | CCL6 | P27784 | http://www.uniprot.org/uniprot/P27784 |
| CD27 | CD27 | P41272 | http://www.uniprot.org/uniprot/P41272 |
| CD27 Ligand | CD70 | P32970 | http://www.uniprot.org/uniprot/P32970 |
| CD30 | CD30 | Q60846 | http://www.uniprot.org/uniprot/Q60846 |
| CD30 Ligand | CD30LG | P32972 | http://www.uniprot.org/uniprot/P32972 |
| CD32a | FCGR2A | P08101 | http://www.uniprot.org/uniprot/P08101 |
| CD36 | CD36 | Q08857 | http://www.uniprot.org/uniprot/Q08857 |
| CD40 | CD40 | Q99NE2 | http://www.uniprot.org/uniprot/Q99NE2 |
| CD40 Ligand | CD40LG | P27548 | http://www.uniprot.org/uniprot/P27548 |
| CD48 | CD48 | Q545K2 | http://www.uniprot.org/uniprot/Q545K2 |
| CD6 | CD6 | Q61003 | http://www.uniprot.org/uniprot/Q61003 |
| CD80 | CD80 | Q00609 | http://www.uniprot.org/uniprot/Q00609 |
| Chemerin | RARRES2 | Q8CHU8 | http://www.uniprot.org/uniprot/Q8CHU8 |
| Chordin | CHRD | Q9Z0E2 | http://www.uniprot.org/uniprot/Q9Z0E2 |
| Clusterin | CLU | Q06890 | http://www.uniprot.org/uniprot/Q06890 |
| CRP | CRP | P14847 | http://www.uniprot.org/uniprot/P14847 |
| CTLA-4 | Ctla4 | Q9QZZ7 | http://www.uniprot.org/uniprot/Q9QZZ7 |
| CXCL16 | CXCL16 | Q5F2D5 | http://www.uniprot.org/uniprot/Q5F2D5 |
| Cystatin C | CST3 | Q544Y0 | http://www.uniprot.org/uniprot/Q544Y0 |
| DAN | PARN | Q8C7N6 | http://www.uniprot.org/uniprot/Q8C7N6 |
| Decorin | DCN | P28654 | http://www.uniprot.org/uniprot/P28654 |
| DKK-1 | DKK1 | Q80UL5 | http://www.uniprot.org/uniprot/Q80UL5 |
| DLL4 | DLL4 | Q9JI71 | http://www.uniprot.org/uniprot/Q9JI71 |
| Dtk | TYRO3 | O09080 | http://www.uniprot.org/uniprot/O09080 |
| E-Cadherin | CDH1 | P09803 | http://www.uniprot.org/uniprot/P09803 |
| EDAR | EDAR | Q9DC43 | http://www.uniprot.org/uniprot/Q9DC43 |
| EGF | EGF | E9QNX6 | http://www.uniprot.org/uniprot/E9QNX6 |
| Endocan | ESM1 | Q9QYY7 | http://www.uniprot.org/uniprot/Q9QYY7 |
| Endoglin | ENG | Q63961 | http://www.uniprot.org/uniprot/Q63961 |
| Eotaxin-1 | CCL11 | P48298 | http://www.uniprot.org/uniprot/P48298 |
| Eotaxin-2 | Ccl24 | Q9JKC0 | http://www.uniprot.org/uniprot/Q9JKC0 |
| Epigen | EPGN | Q8CEX5 | http://www.uniprot.org/uniprot/Q8CEX5 |
| Epiregulin | EREG | Q61521 | http://www.uniprot.org/uniprot/Q61521 |
| E-Selectin | SELE | Q00690 | http://www.uniprot.org/uniprot/Q00690 |
| Fas | Fas | Q6GT31 | http://www.uniprot.org/uniprot/Q6GT31 |
| Fas Ligand | FASLG | P41047 | http://www.uniprot.org/uniprot/P41047 |
| Fetuin A | AHSG | P29699 | http://www.uniprot.org/uniprot/P29699 |
| Flt-3 Ligand | FLT3LG | P49772 | http://www.uniprot.org/uniprot/P49772 |
| Fractalkine | CX3CL1 | O35188 | http://www.uniprot.org/uniprot/O35188 |
| Galectin-1 | LGALS1 | P17601 | http://www.uniprot.org/uniprot/P17601 |
| Galectin-3 | LGALS3 | Q8C253 | http://www.uniprot.org/uniprot/Q8C253 |
| Galectin-7 | LGALS7 | Q9CRB1 | http://www.uniprot.org/uniprot/Q9CRB1 |
| Gas 1 | GAS1 | Q01721 | http://www.uniprot.org/uniprot/Q01721 |
| Gas 6 | Gas6 | Q99K57 | http://www.uniprot.org/uniprot/Q99K57 |
| GCSF | CSF3 | P09920 | http://www.uniprot.org/uniprot/P09920 |
| GITR | TNFRSF18 | Q9JKR2 | http://www.uniprot.org/uniprot/Q9JKR2 |
| GITR Ligand | TNFSF18 | Q7TS55 | http://www.uniprot.org/uniprot/Q7TS55 |
| GM-CSF | CSF2 | P01587 | http://www.uniprot.org/uniprot/P01587 |
| gp130 | Il6st | Q00560 | http://www.uniprot.org/uniprot/Q00560 |
| Granzyme B | GZMB | Q3V2P1 | http://www.uniprot.org/uniprot/Q3V2P1 |
| Gremlin-1 | GREM1 | O70326 | http://www.uniprot.org/uniprot/O70326 |
| H60 | H60a | Q3TDZ7 | http://www.uniprot.org/uniprot/Q3TDZ7 |
| HAI-1 | SPINT1 | Q9R097 | http://www.uniprot.org/uniprot/Q9R097 |
| HGF | HGF | Q53WS5 | http://www.uniprot.org/uniprot/Q53WS5 |
| HGFR | MET | D3YVY2 | http://www.uniprot.org/uniprot/D3YVY2 |
| I-309 | CCL1 | P14098 | http://www.uniprot.org/uniprot/P14098 |
| ICAM-1 | ICAM1 | P13597 | http://www.uniprot.org/uniprot/P13597 |
| IFN-gamma | IFNG | P01580 | http://www.uniprot.org/uniprot/P01580 |
| IFN-gamma R1 | IFNGR1 | P15261 | http://www.uniprot.org/uniprot/P15261 |
| IGF-1 | IGF1 | P05018 | http://www.uniprot.org/uniprot/P05018 |
| IGFBP-2 | IGFBP2 | P47877 | http://www.uniprot.org/uniprot/P47877 |
| IGFBP-3 | IGFBP3 | P47878 | http://www.uniprot.org/uniprot/P47878 |
| IGFBP-5 | IGFBP5 | Q07079 | http://www.uniprot.org/uniprot/Q07079 |
| IGFBP-6 | IGFBP6 | Q91X24 | http://www.uniprot.org/uniprot/Q91X24 |
| IL-1 alpha | IL1A | P01582 | http://www.uniprot.org/uniprot/P01582 |
| IL-1 beta | IL1B | P10749 | http://www.uniprot.org/uniprot/P10749 |
| IL-1 R4 | IL1RL1 | Q05208 | http://www.uniprot.org/uniprot/Q05208 |
| IL-1 ra | IL1RN | P25085 | http://www.uniprot.org/uniprot/P25085 |
| IL-10 | IL10 | Q0VBJ1 | http://www.uniprot.org/uniprot/Q0VBJ1 |
| IL-12 p40 | IL12B | P43432 | http://www.uniprot.org/uniprot/P43432 |
| IL-12 p70 | IL12A | P43431 | http://www.uniprot.org/uniprot/P43431 |
| IL-13 | IL13 | P20109 | http://www.uniprot.org/uniprot/P20109 |
| IL-15 | IL15 | Q1AHQ7 | http://www.uniprot.org/uniprot/Q1AHQ7 |
| IL-17 RB | IL17RB | Q9JIP3 | http://www.uniprot.org/uniprot/Q9JIP3 |
| IL-17A | IL17A | Q60971 | http://www.uniprot.org/uniprot/Q60971 |
| IL-17B | IL17B | Q99MY3 | http://www.uniprot.org/uniprot/Q99MY3 |
| IL-17E | IL25 | Q8VHH8 | http://www.uniprot.org/uniprot/Q8VHH8 |
| IL-17F | IL17F | Q7TNI7 | http://www.uniprot.org/uniprot/Q7TNI7 |
| IL-2 | IL2 | Q791T3 | http://www.uniprot.org/uniprot/Q791T3 |
| IL-2 R alpha | IL2RA | Q61731 | http://www.uniprot.org/uniprot/Q61731 |
| IL-20 | IL20 | Q9JKV9 | http://www.uniprot.org/uniprot/Q9JKV9 |
| IL-21 | IL21 | E9PX58 | http://www.uniprot.org/uniprot/E9PX58 |
| IL-22 | IL22 | Q14BB3 | http://www.uniprot.org/uniprot/Q14BB3 |
| IL-23 p19 | IL23A | Q9EQ14 | http://www.uniprot.org/uniprot/Q9EQ14 |
| IL-28A | IFNL2 | Q4VK74 | http://www.uniprot.org/uniprot/Q4VK74 |
| IL-3 | IL3 | P01586 | http://www.uniprot.org/uniprot/P01586 |
| IL-3 R beta | Csf2rb | P26955 | http://www.uniprot.org/uniprot/P26955 |
| IL-33 | IL33 | Q8BVZ5 | http://www.uniprot.org/uniprot/Q8BVZ5 |
| IL-4 | IL4 | P07750 | http://www.uniprot.org/uniprot/P07750 |
| IL-5 | IL5 | P04401 | http://www.uniprot.org/uniprot/P04401 |
| IL-6 | IL6 | P08505 | http://www.uniprot.org/uniprot/P08505 |
| IL-7 | IL7 | P10168 | http://www.uniprot.org/uniprot/P10168 |
| IL-7 R alpha | IL7R | Q9R0C1 | http://www.uniprot.org/uniprot/Q9R0C1 |
| IL-9 | IL9 | P15247 | http://www.uniprot.org/uniprot/P15247 |
| I-TAC | CXCL11 | Q9JHH5 | http://www.uniprot.org/uniprot/Q9JHH5 |
| JAM-A | F11R | Q8VC39 | http://www.uniprot.org/uniprot/Q8VC39 |
| KC | CXCL1 | P12850 | http://www.uniprot.org/uniprot/P12850 |
| Kremen-1 | KREMEN1 | Q99N43 | http://www.uniprot.org/uniprot/Q99N43 |
| Leptin | LEP | P41160 | http://www.uniprot.org/uniprot/P41160 |
| Leptin R | LEPR | Q61215 | http://www.uniprot.org/uniprot/Q61215 |
| Limitin | Limitin | Q9R1T0 | <https://www.uniprot.org/uniprot/Q9R1T0> |
| Lipocalin-2 | LCN2 | Q3UE34 | http://www.uniprot.org/uniprot/Q3UE34 |
| LIX | Cxcl5 | P50228 | http://www.uniprot.org/uniprot/P50228 |
| LOX-1 | OLR1 | Q9EQ09 | http://www.uniprot.org/uniprot/Q9EQ09 |
| L-Selectin | SELL | Q3TCF3 | http://www.uniprot.org/uniprot/Q3TCF3 |
| Lungkine | CXCL15 | Q3UQ15 | http://www.uniprot.org/uniprot/Q3UQ15 |
| Lymphotactin | XCL1 | P47993 | http://www.uniprot.org/uniprot/P47993 |
| MAdCAM-1 | MADCAM1 | Q61278 | http://www.uniprot.org/uniprot/Q61278 |
| Marapsin | PRSS27 | Q14A25 | http://www.uniprot.org/uniprot/Q14A25 |
| MBL-2 | MBL2 | P41317 | http://www.uniprot.org/uniprot/P41317 |
| MCP-1 | CCL2 | Q9QYD7 | http://www.uniprot.org/uniprot/Q9QYD7 |
| MCP-5 | Ccl12 | Q9QYD6 | http://www.uniprot.org/uniprot/Q9QYD6 |
| M-CSF | CSF1 | P07141 | http://www.uniprot.org/uniprot/P07141 |
| MDC | CCL22 | O88430 | http://www.uniprot.org/uniprot/O88430 |
| Meteorin | METRN | Q8CI64 | http://www.uniprot.org/uniprot/Q8CI64 |
| MFG-E8 | Mfge8 | Q9R1X9 | http://www.uniprot.org/uniprot/Q9R1X9 |
| MIG | CXCL9 | P18340 | http://www.uniprot.org/uniprot/P18340 |
| MIP-1 alpha | CCL3 | P14096 | http://www.uniprot.org/uniprot/P14096 |
| MIP-1 beta | CCL4 | P14097 | http://www.uniprot.org/uniprot/P14097 |
| MIP-1 gamma | Ccl9 | Q5QNW2 | http://www.uniprot.org/uniprot/Q5QNW2 |
| MIP-2 | Cxcl2 | P10889 | http://www.uniprot.org/uniprot/P10889 |
| MIP-3 alpha | CCL20 | O89093 | http://www.uniprot.org/uniprot/O89093 |
| MIP-3 beta | CCL19 | O70460 | http://www.uniprot.org/uniprot/O70460 |
| MMP-10 | MMP10 | O55123 | http://www.uniprot.org/uniprot/O55123 |
| MMP-2 | MMP2 | P33434 | http://www.uniprot.org/uniprot/P33434 |
| MMP-3 | MMP3 | P28862 | http://www.uniprot.org/uniprot/P28862 |
| Neprilysin | MME | Q6NXX5 | http://www.uniprot.org/uniprot/Q6NXX5 |
| Nope | IGDCC4 | Q9EQS9 | http://www.uniprot.org/uniprot/Q9EQS9 |
| NOV | NOV | Q64299 | http://www.uniprot.org/uniprot/Q64299 |
| Osteoactivin | GPNMB | Q99P91 | http://www.uniprot.org/uniprot/Q99P91 |
| Osteopontin | SPP1 | P19008 | http://www.uniprot.org/uniprot/P19008 |
| Osteoprotegerin | TNFRSF11B | O70202 | http://www.uniprot.org/uniprot/O70202 |
| OX40 Ligand | TNFSF4 | P43488 | <https://www.uniprot.org/uniprot/P43488> |
| P-Cadherin | CDH3 | Q8BSL6 | http://www.uniprot.org/uniprot/Q8BSL6 |
| PDGF-AA | PDGFA | P20033 | http://www.uniprot.org/uniprot/P20033 |
| Pentraxin-3 | PTX3 | P48759 | http://www.uniprot.org/uniprot/P48759 |
| Periostin | POSTN | Q62009 | http://www.uniprot.org/uniprot/Q62009 |
| Persephin | PSPN | A1L3Q1 | http://www.uniprot.org/uniprot/A1L3Q1 |
| Platelet Factor 4 | CXCL14 | Q9WUQ5 | http://www.uniprot.org/uniprot/Q9WUQ5 |
| PlGF-2 | Pgf | P49764 | http://www.uniprot.org/uniprot/P49764 |
| Progranulin | GRN | P28798 | http://www.uniprot.org/uniprot/P28798 |
| Prolactin | PRL | P06879 | http://www.uniprot.org/uniprot/P06879 |
| Pro-MMP-9 | Mmp9 | Q80XI8 | http://www.uniprot.org/uniprot/Q80XI8 |
| Prostasin | PRSS8 | Q99L44 | http://www.uniprot.org/uniprot/Q99L44 |
| P-Selectin | SELP | Q01102 | http://www.uniprot.org/uniprot/Q01102 |
| RAGE | MOK | Q9WVS4 | http://www.uniprot.org/uniprot/Q9WVS4 |
| RANTES | CCL5 | P30882 | http://www.uniprot.org/uniprot/P30882 |
| Renin 1 | REN | P06281 | http://www.uniprot.org/uniprot/P06281 |
| Resistin | RETN | Q99P87 | http://www.uniprot.org/uniprot/Q99P87 |
| SCF | KITLG | P20826 | http://www.uniprot.org/uniprot/P20826 |
| SDF-1 alpha | CXCL12 | P40224 | http://www.uniprot.org/uniprot/P40224 |
| sFRP-3 | FRZB | O09075 | http://www.uniprot.org/uniprot/O09075 |
| SLAM | SLAMF1 | Q9QXZ3 | http://www.uniprot.org/uniprot/Q9QXZ3 |
| Sonic Hedgehog N-Terminal | SHH | Q62226 | <https://www.uniprot.org/uniprot/Q62226> |
| TACI | TNFRSF13B | Q9DBZ3 | http://www.uniprot.org/uniprot/Q9DBZ3 |
| TARC | CCL17 | Q9WUZ6 | http://www.uniprot.org/uniprot/Q9WUZ6 |
| TCK-1 | Ppbp | Q9EQI5 | http://www.uniprot.org/uniprot/Q9EQI5 |
| TECK | CCL25 | O35903 | http://www.uniprot.org/uniprot/O35903 |
| Testican 3 | SPOCK3 | Q8BKV0 | http://www.uniprot.org/uniprot/Q8BKV0 |
| TGF beta 1 | TGFB1 | P04202 | http://www.uniprot.org/uniprot/P04202 |
| Thrombopoietin | THPO | P40226 | http://www.uniprot.org/uniprot/P40226 |
| TIM-1 | HAVCR1 | Q8VIM1 | http://www.uniprot.org/uniprot/Q8VIM1 |
| TNF alpha | TNF | Q62326 | http://www.uniprot.org/uniprot/Q62326 |
| TNF RI | TNFRSF1A | P25118 | http://www.uniprot.org/uniprot/P25118 |
| TNF RII | TNFRSF1B | O88734 | http://www.uniprot.org/uniprot/O88734 |
| TRAIL | TNFSF10 | P50592 | http://www.uniprot.org/uniprot/P50592 |
| Trance | TNFSF11 | Q9R1Y0 | http://www.uniprot.org/uniprot/Q9R1Y0 |
| TREM-1 | TREM1 | Q3TAL7 | http://www.uniprot.org/uniprot/Q3TAL7 |
| TREML1 | TREML1 | Q8K558 | http://www.uniprot.org/uniprot/Q8K558 |
| TROY | TNFRSF19 | Q80T13 | http://www.uniprot.org/uniprot/Q80T13 |
| Tryptase epsilon | PRSS22 | Q9ER10 | http://www.uniprot.org/uniprot/Q9ER10 |
| TSLP | TSLP | Q9JIE6 | http://www.uniprot.org/uniprot/Q9JIE6 |
| TWEAK | TNFSF12 | O54907 | http://www.uniprot.org/uniprot/O54907 |
| TWEAK R | TNFRSF12A | Q9CR75 | http://www.uniprot.org/uniprot/Q9CR75 |
| VCAM-1 | VCAM1 | P29533 | http://www.uniprot.org/uniprot/P29533 |
| VEGF-A | VEGFA | Q00731 | http://www.uniprot.org/uniprot/Q00731 |
| VEGF-B | VEGFB | P49766 | <https://www.uniprot.org/uniprot/P49766> |
| VEGF-D | FIGF | A2AIH3 | http://www.uniprot.org/uniprot/A2AIH3 |
| VEGFR1 | Flt1 | Q61517 | http://www.uniprot.org/uniprot/Q61517 |
| VEGFR2 | KDR | Q3UQZ6 | http://www.uniprot.org/uniprot/Q3UQZ6 |
| VEGFR3 | FLT4 | P35917 | http://www.uniprot.org/uniprot/P35917 |

**Supplemental Figure 1. Secretome Heatmap.** Large heatmap and dendrogram of complete microarray panel. Provided to facilitate readability of heatmap (Fig. 2).

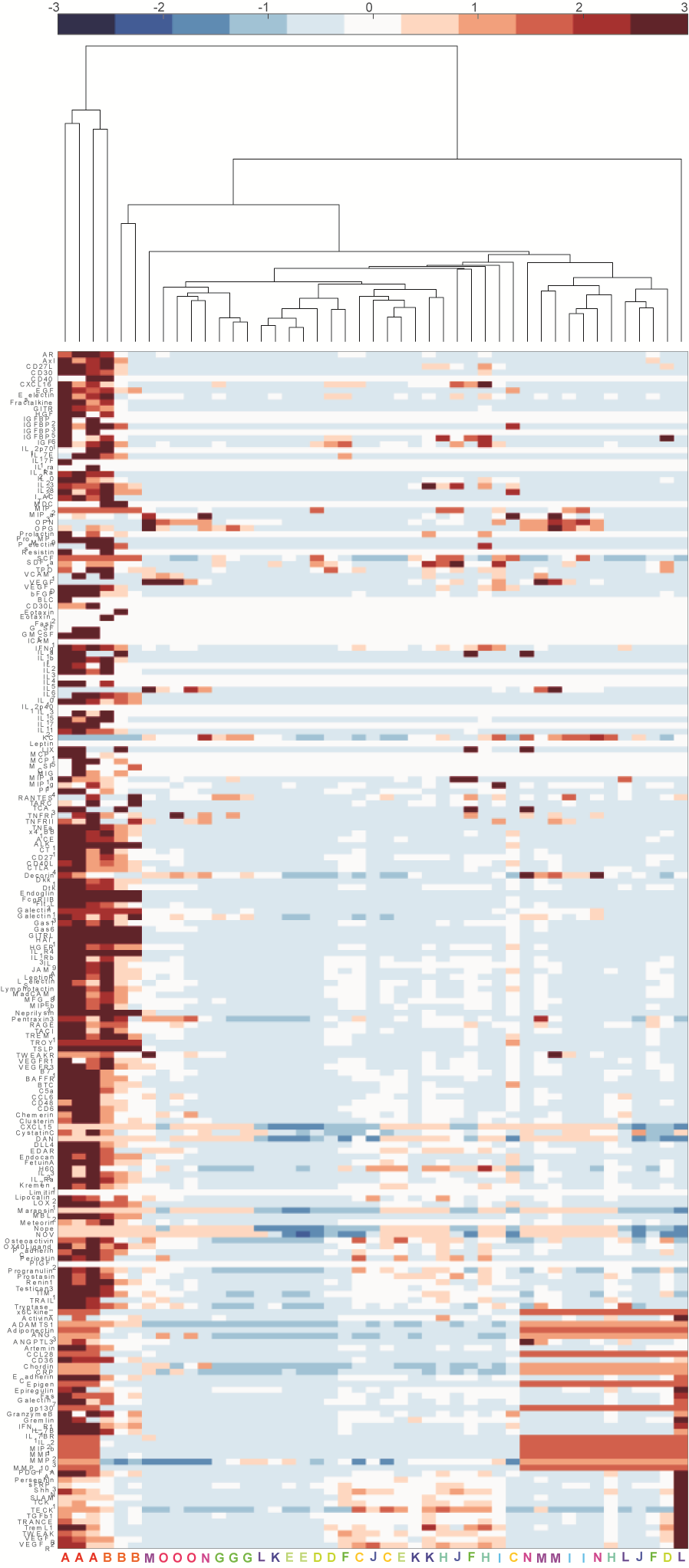

**Supplemental Figure 2. Cell-cycle Proportions.** Analysis of cell-cycle status (G0, G1, SGM) for both HSPC only (Single culture; 1:0 HSPC:MSPC) and 1:1 HSPC:MSPC co-culture (Co-Culture) in response to exogenous cytokines (TGF-β1, latent MMP3, activated MMP3, TROY, cRP) or combinations of all factors (using either latent or activated MMP3).
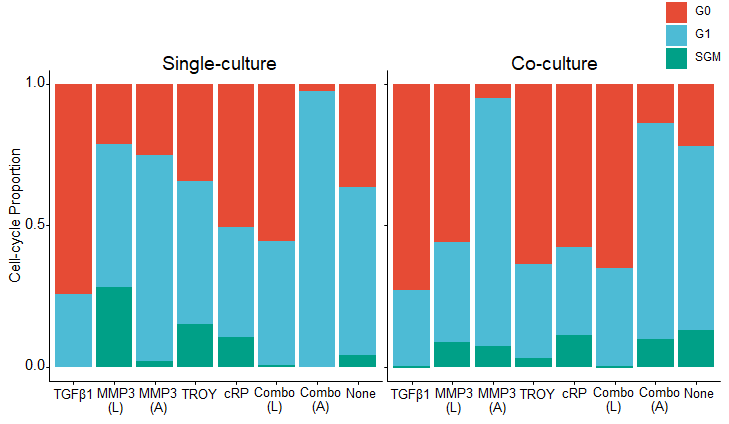

**Supplemental Figure 3. STRING Protein-Interaction.** Interactions of each of the 15 cytokines identified by iterative filter method PLSR using STRING-db.org.

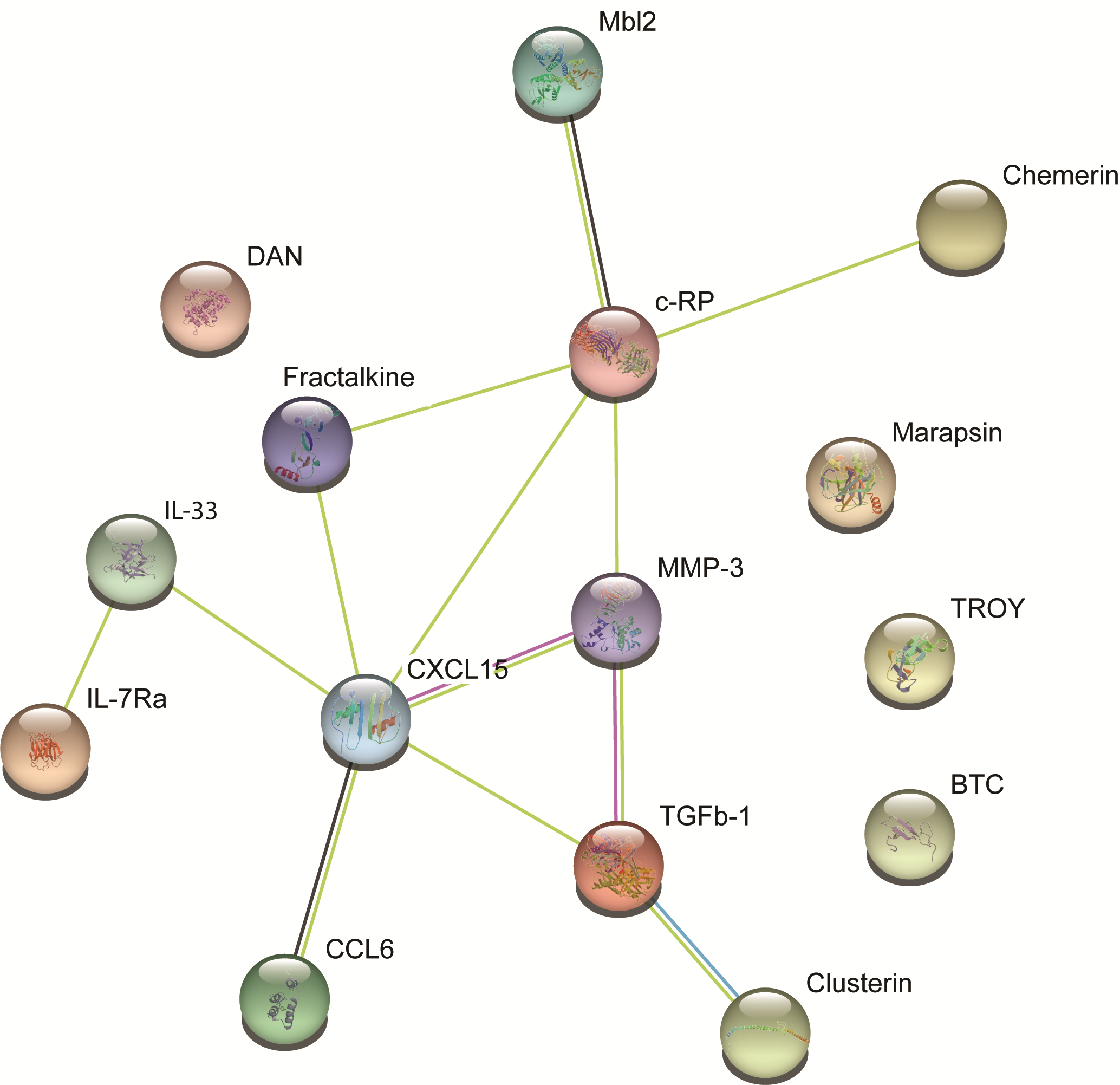
